# Associative learning drives longitudinally-graded presynaptic plasticity of neurotransmitter release along axonal compartments

**DOI:** 10.1101/2021.06.08.447536

**Authors:** Aaron Stahl, Nathaniel C. Noyes, Tamara Boto, Miao Jing, Jianzhi Zeng, Lanikea B. King, Yulong Li, Ronald L. Davis, Seth M. Tomchik

## Abstract

Anatomical and physiological compartmentalization of neurons is a mechanism to increase the computational capacity of a circuit, and a major question is what role axonal compartmentalization plays. Axonal compartmentalization may enable localized, presynaptic plasticity to alter neuronal output in a flexible, experience-dependent manner. Here we show that olfactory learning generates compartmentalized, bidirectional plasticity of acetylcholine release that varies across the longitudinal compartments of *Drosophila* mushroom body (MB) axons. The directionality of the learning-induced plasticity depends on the valence of the learning event (aversive vs. appetitive), varies linearly across proximal to distal compartments following appetitive conditioning, and correlates with learning-induced changes in downstream mushroom body output neurons (MBONs) that modulate behavioral action selection. Potentiation of acetylcholine release was dependent on the Ca_V_2.1 calcium channel subunit *cacophony*. In addition, contrast between the positive conditioned stimulus and other odors required the inositol triphosphate receptor (IP_3_R), which was required to maintain responsivity to odors in untrained conditions. Downstream from the mushroom body, a set of MBONs that receive their input from the γ3 MB compartment were required for normal appetitive learning, suggesting that they represent a key node through which discriminative effects influence appetitive memory and decision-making. These data demonstrate that learning drives valence-correlated, compartmentalized, bidirectional potentiation and depression of synaptic neurotransmitter release, which rely on distinct mechanisms and are distributed across axonal compartments in a learning circuit.

## Introduction

Neuronal dendrites carry out computations through compartmentalized signaling, while axons have long been considered to carry signals to their terminal fields relatively uniformly following spike initiation. However, anatomical and physiological compartmentalization of axons has been recently documented in neurons from worms through mammals (Boto et al., 2014; Cohn et al., 2015; Hendricks et al., 2012; Rowan et al., 2016). How axonal compartmentalization influences information flow across neuronal circuits and modulates behavioral outcomes is not understood. One functional role for axonal compartmentalization may be to enable localized, presynaptic plasticity to alter output from select axon compartments in a flexible, experience-dependent manner. This would vastly enhance the neuron’s flexibility and computational capabilities. One potential function of such compartmentalization would allow independent modulation of axonal segments and/or synaptic release sites by biologically-salient events, such as sensory stimuli that drive learning.

The anatomical organization of the *Drosophila* mushroom body (MB) makes it an exemplary testbed to study how sensory information is processed during learning and rerouted to alter behavioral outcomes. The MB encodes odor in sparse representations across intrinsic MB neurons, which are arranged in several parallel sets. They project axons in fasciculated bundles into several anatomically-distinct, but spatially adjacent lobes (α/β, α′/ β′, and γ) (Crittenden et al., 1998). These bundled axons are longitudinally subdivided into discrete tiled compartments (Aso et al., 2014a). Each compartment receives afferent neuromodulatory input from unique dopaminergic neurons (Aso et al., 2014a; Mao and Davis, 2009), and innervates unique efferent mushroom body output neurons (MBONs) (Aso et al., 2014a). Each set of dopaminergic neurons plays an individual role in learning, with some conveying aversive teaching signals (Schroll et al., 2006; Schwaerzel et al., 2003), others conveying positive teaching signals (Liu et al., 2012; Yamagata et al., 2015), and a third class modulating memory strength without driving valence (Boto et al., 2019). Likewise, each MBON has a unique effect on behavioral approach and avoidance, with some biasing the animal to approach, others biasing the animal to avoidance, and some having no effect (Aso et al., 2014b; Perisse et al., 2016; Placais et al., 2013; Sejourne et al., 2011).

A major question in learning and memory is how presynaptic plasticity contributes to reweight the flow of sensory signals down each of the downstream “approach” or “avoidance” circuits, altering action selection and memory retrieval. In naïve conditions, *Drosophila* dopaminergic circuits modulate cAMP in a compartmentalized fashion along the MB axons (Boto et al., 2014). This compartmentalized dopaminergic signaling can independently modulate Ca^2+^ responses in each compartment, as well as the responses of the downstream valence-coding MBONs (Cohn et al., 2015). Dopamine-dependent heterosynaptic depression at the MB-MBON synapse modulates learning (Hige et al., 2015a). Therefore, presynaptic plasticity in the MB neurons within each compartment could theoretically drive the changes in MBON responsiveness that guide behavioral learning (Zhang et al., 2019). However, manipulation of the “aversive” protocerebral posterior lateral 1 (PPL1) dopaminergic neurons does not detectably alter Ca^2+^ signals in MB neurons (Boto et al., 2019; Hige et al., 2015a). Furthermore, Ca^2+^ responses in MB neurons are uniformly potentiated across compartments with appetitive classical conditioning protocols and unaltered in MB neurons following aversive protocols (Louis et al., 2018). This raises the question of how local, compartmentalized synaptic plasticity in MB neurons drives coherent changes in downstream MBONs to modulate action selection during memory retrieval. Learning/dopamine-induced plasticity has been demonstrated in the downstream MBONs (Berry et al., 2018; Hige et al., 2015a; Hige et al., 2015b; Owald et al., 2015), with dopamine also acting directly on MBONs (Takemura et al., 2017). Feedforward inhibition among MBONs that drive opposing behavioral outcomes provides a mechanism explaining how valence coding in MBONs could be generated (Perisse et al., 2016). Yet this does not explain the compartmentalized, dopamine-dependent plasticity in MB neurons themselves or the necessity for dopamine receptors and downstream signaling molecules in the intrinsic MB neurons (Kim et al., 2007; McGuire et al., 2003; Zars et al., 2000).

Here we describe how learning alters the flow of information through the MB via synaptic release of the putative MB neurotransmitter (Barnstedt et al., 2016), using genetically-encoded indicators of synaptic acetylcholine neurotransmission. The data reveal that learning alters the compartmentalized axonal acetylcholine release from *Drosophila* mushroom body (MB) neurons in valence-specific spatiotemporal patterns, via distinct molecular mechanisms, driving behavioral alterations via modulation of specific downstream output neurons.

## Results

### Associative learning modulates neurotransmitter release in a spatially-distinct manner across longitudinal axonal compartments

Synapses within each MB compartment transmit olfactory information from MB neurons to compartment-specific MBONs (Fig. 1A, 5A,B) (Aso et al., 2014a; Tanaka et al., 2008). The MBONs exert distinct and often-opposing effects on behavior, with some innately promoting approach and others promoting avoidance (Aso et al., 2014b; Berry et al., 2018; Ichinose et al., 2015; Owald et al., 2015; Perisse et al., 2016; Placais et al., 2013; Sayin et al., 2019; Sejourne et al., 2011). Synaptic depression has been observed in the MB-MBON synapses following pairing of odor with stimulation of PPL1 neurons that are critical for aversive learning (Hige et al., 2015a), suggesting that depression may be a primary mechanism for learning at these synapses (Barnstedt et al., 2016; Cohn et al., 2015; Handler et al., 2019; Owald et al., 2015; Perisse et al., 2016; Sejourne et al., 2011). One synapse downstream, some MBONs exhibit bidirectional responses to conditioning, though the major described mechanism involves a sign change that occurs postsynaptic to the MBs (polysynaptic feedforward inhibition) (Owald et al., 2015; Perisse et al., 2016). To test for the presence, directionality, and variation of presynaptic plasticity across MB axonal compartments, we expressed a synaptic ACh sensor to monitor neurotransmitter release from MB neurons *in vivo* (Zhang et al., 2019). The genetically-encoded ACh reporter, GPCR-Activation–Based-ACh sensor (GRAB-ACh) (Fig. 1A) (Jing et al., 2019; Jing et al., 2018; Zhang et al., 2019), was expressed in MB neurons using the 238Y-Gal4 driver. Appetitive conditioning was carried out, monitoring ACh release from the γ lobe compartments evoked by the CS+ and CS− before and after pairing odor with sucrose (Fig. S1). Responses were compared to those in odor-only control cohorts to determine whether any learning-induced changes resulted from potentiation or depression. We quantified several parameters (Fig. S1), including how the responses changed after conditioning (the within-treatment post/pre). In addition, we compared the CS+ and CS− responses after conditioning (CS:CS−), which mimics the putative comparison the animal makes during associative memory retrieval. Finally, we compared the change in CS+ and CS−, Δ(post/pre), to their respective odor-only controls to quantify whether they were potentiated or depressed by conditioning, accounting for any sensory adaptation (Figs. 1 F-J, S2).

**Figure 1.**
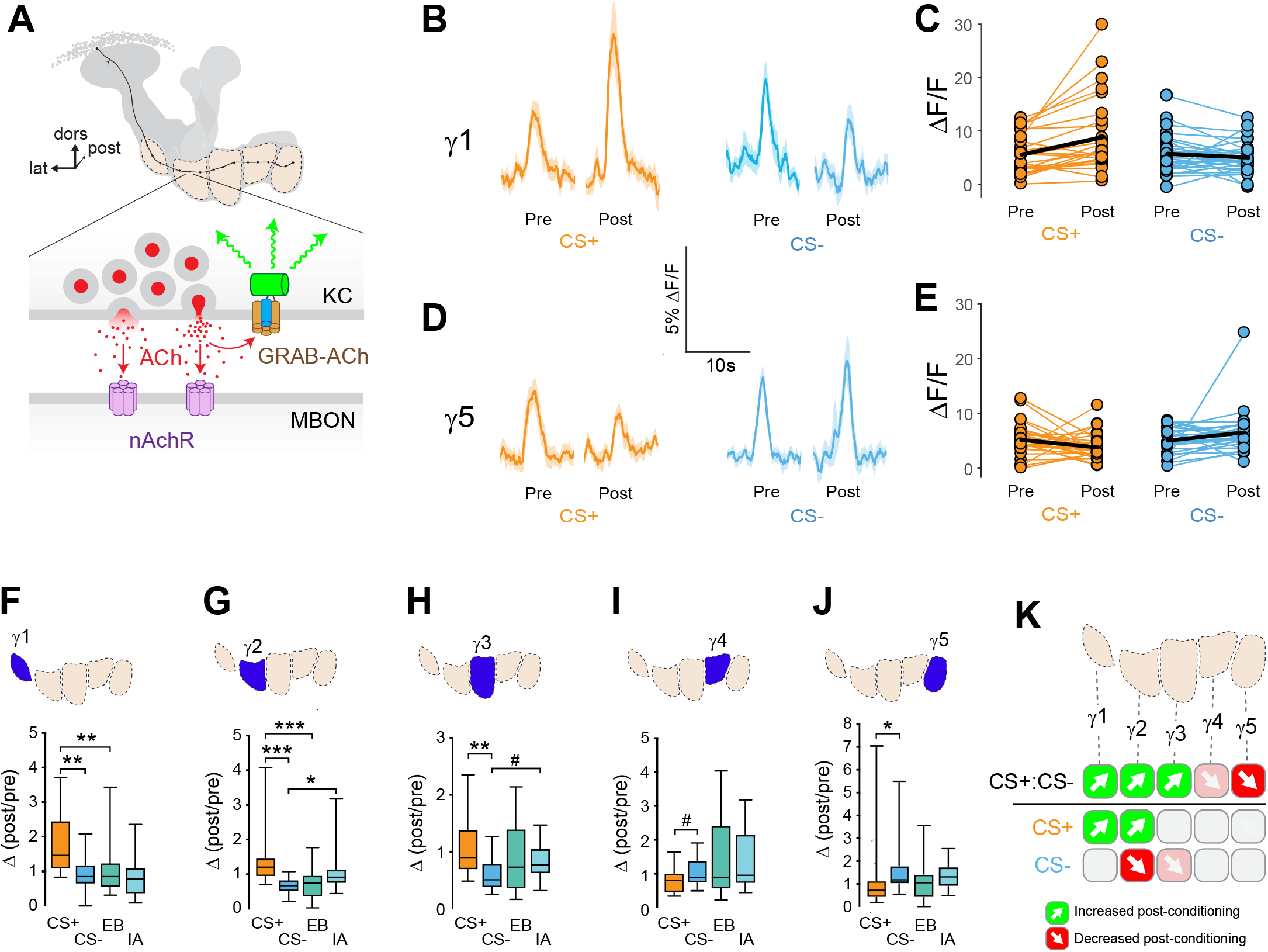
Compartment-specific alterations of ACh release in the MB following appetitive conditioning. **(A)** Diagram of the GRAB-ACh reporter expressed in presynaptic terminals of MB neurons (Kenyon cells: MB). nAChR: nicotinic acetylcholine receptor; dors: dorsal; lat: lateral, post: posterior; MBON: mushroom body output neuron. **(B)** Time series traces showing odor-evoked GRAB-ACh responses pre- and post-conditioning. Responses were imaged to both the CS+ (ethyl butyrate: EB) and CS− (isoamyl acetate: IA) odor in the γ1 compartment, and the line and shading represent the mean ± SEM. **(C)** Quantification of the pre- and post-conditioning responses to the CS+ (EB) and CS− (IA) from the γ1 compartment from individual animals (n = 27), with the mean graphed as a black line. **(D)** Time series traces imaged from the γ5 compartment, graphed as in panel B. **(E)** Quantification of peak responses from the γ5 compartment, graphed as in panel C. **(F-J)** Change in odor-evoked responses (Post/pre responses), following conditioning (CS+ and CS−) or odor-only presentation (EB and IA). *p<0.01, **p<0.001, ***p<0.0001; n = 27 (Kruskal-Wallis/Bonferonni). **(F)** γ1compartment. **(G)** γ2 compartment. **(H)** γ3 compartment. #p = 0.0169. **(I)** γ4 compartment. #p = 0.0868. (J) 5 compartment. **(K)** Summary of plasticity in ACh release across lobe compartments. Green up arrows indicate increases in the CS+:CS− (1st row) or potentiation of the CS+ response (relative to odor-only controls; 2nd row), while red down arrows indicate decreases in the CS+:CS− (1st row) or depression of the CS− (relative to odor-only controls; 3rd row).

Appetitive conditioning produced plasticity in ACh release that varied across the axonal compartments of the MB γ lobe in several key ways (Fig. 1). Conditioning significantly increased CS+ responses relative to the CS− responses (↑CS+:CS−) in the three most proximal γ lobe compartments: γ1, γ2, and γ3 (Figs. 1 B,C,F-H; S2). In each of these compartments, this was due to different underlying mechanisms. In the γ1 compartment, the CS+ response was potentiated; i.e., following appetitive conditioning, the Δ (post/pre) CS+ response was larger than the ethyl butyrate (EB) odor-only control, while the CS− did not significantly differ from the isoamyl acetate (IA) odor-only control (Figs. 1F, S2). In the γ2 compartment, both the CS+ was potentiated and the CS− depressed relative to the odor-only controls (Figs. 1G, S2). Finally, in the γ3 compartment, neither was significantly altered relative to odor-only controls at the Bonferroni-corrected α=0.01 level, but there was a strong trend toward depression with the CS− group (Figs. 1H, S2). Thus, there was a spatial gradient of CS+ potentiation in γ1, shifting from CS+ potentiation in γ1 toward CS− depression in γ3, with the spatially-intermediate γ2 exhibiting both. This gradient of CS+:CS− plasticity suggests that both the CS+ and CS− contribute to learning by modulating MB output.

In the distal γ4-γ5 compartments, appetitive conditioning produced plasticity in the opposite direction to that in the proximal γ compartments. In these compartments, the CS+ response was reduced relative to the CS− (↓CS+:CS−) (Figs. 1D, E, I-J, S2). This effect was significant in the γ5 compartment (Fig. 1J), while γ4 exhibited a trend in the same direction (Fig. 1I). In these compartments, the effect could not be unambiguously assigned to CS+ depression, though there was no evidence of CS− potentiation (Figs. 1 I,J, S2). Overall, appetitive conditioning produced net enhancement of CS+ responsivity in γ1-γ3 compartments, which was derived from a proximal-to-distal gradient of CS+ potentiation to CS− depression, and net reduction of CS+ responsivity in γ4-γ5 (Fig. 1K). Thus, the plasticity was bidirectional between the proximal and distal axonal compartments. This likely contributes to approach behavior by simultaneously enhancing the conditioned odor-evoked activation of downstream “approach” circuits and inhibition of “avoidance” MBON circuits.

### Conditioning with opposing valence stimuli generates bidirectional presynaptic plasticity within axonal compartments

The above data suggested that appetitive conditioning produced synaptic potentiation in the proximal γ lobe compartments. Yet synaptic depression is the main described plasticity mechanism at the MB-MBON synapses following olfactory conditioning (Barnstedt et al., 2016; Modi et al., 2020; Owald et al., 2015; Perisse et al., 2016; Sejourne et al., 2011; Zhang and Roman, 2013; Zhang et al., 2019). In the γ1 compartment, where it has been examined in detail with electrophysiology, aversive reinforcement substitution produces synaptic depression (Hige et al., 2015a). Since many of these studies involved aversive conditioning, we reasoned that appetitive and aversive conditioning may produce bidirectional plasticity, with the sign/directionality matching postsynaptic MBON valence. To test this, we examined whether aversive conditioning produced the opposite effect in the same compartments as appetitive conditioning had. ACh release from MB neurons was imaged with GRAB-ACh and flies were trained with an aversive odor-shock conditioning protocol (Fig. 2A). In these experiments, we focused on the γ2-γ5 compartments, as the fly was mounted at a higher angle, making the GRAB-ACh signal difficult to simultaneously visualize from γ1 along with that of the other compartments. Following aversive conditioning, there was a reduction in the CS+ response relative to the CS− (↓CS+:CS−) in the γ2 and γ3 compartments (Fig. 2 C-H). This was due to depression in the CS+ response, as the post-conditioning CS+ response was significantly smaller than odor-only controls. The γ4 and γ5 compartments exhibited no significant change in ACh release (Fig 2 I-J). When compared to appetitive conditioning, aversive stimuli produced plasticity that created a sign flip in the γ2 and γ3 compartments (Figs. 1K, 2K). Thus, appetitive and aversive conditioning produced bidirectional plasticity across multiple compartments, which was due to localized plasticity within MB lobe. The aversive conditioning-induced depression likely represents a presynaptic contribution to learning-induced changes in odor responsivity among postsynaptic MBONs (Berry et al., 2018; Hige et al., 2015a; Owald et al., 2015; Zhang et al., 2019).

**Figure 2.**
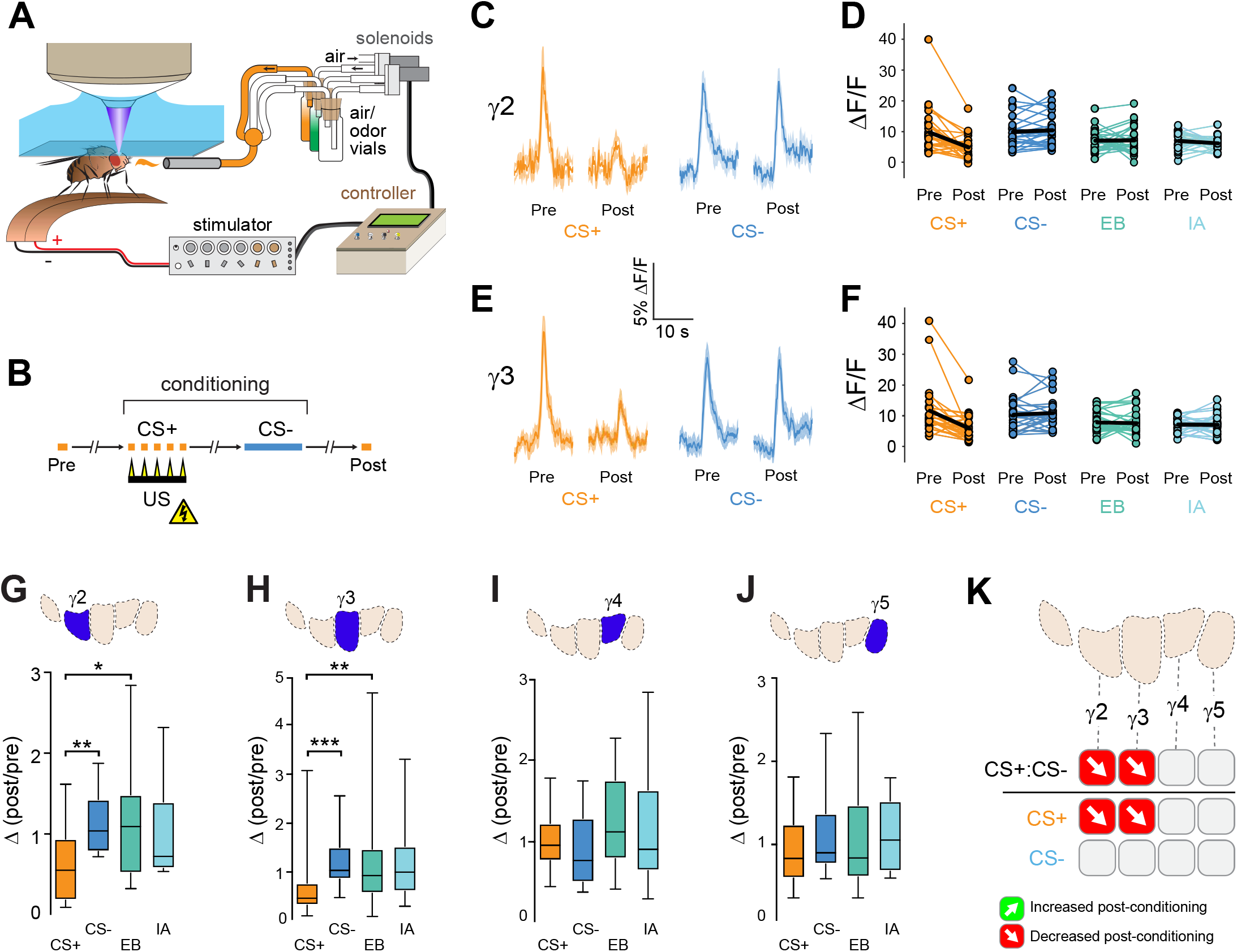
Compartment-specific alterations of ACh release in the MB following aversive conditioning. **(A)** Diagram of the aversive conditioning apparatus. **(B)** Aversive conditioning experimental protocol, pairing an odor (the CS+) with an electric shock unconditioned stimulus (US) (6 shocks, 60V). A second odor, the CS− was presented 5 min after pairing the CS+ and US. One odor was imaged before (Pre) and after (Post) conditioning per animal (CS+ diagrammed here). **(C)** Time series traces showing odor-evoked GRAB-ACh responses pre- and post-conditioning. Responses were imaged to both the CS+ (ethyl butyrate: EB) and CS− (isoamyl acetate: IA) odor in the γ2 compartment, and the line and shading represent the mean ± SEM. **(D)** Quantification of the peak pre- and post-conditioning responses to the CS+ (EB) and CS− (IA) from the γ2compartment from individual animals (n = 27), with the mean graphed as a black line. **(E)** Time series traces imaged from the γ3 compartment, graphed as in panel B. **(F)** Quantification of peak responses from the γ3 compartment, graphed as in panel C. **(G-J)** Change in odor-evoked responses (Post/pre responses), following conditioning (CS+ and CS−) or odor-only presentation (EB and IA). *p<0.01, **p<0.001, ***p<0.0001; n = 27 (Kruskal-Wallis/Bonferonni). **(G)** γ2 compartment. **(H)** γ3 compartment. **(I)** γ4 compartment. **(J)** γ5 compartment. **(K)** Summary of plasticity in ACh release across γ lobe compartments. Red down arrows indicate decreases in the CS+:CS− (1st row) or depression of the CS+ (relative to odor-only controls; 2^nd^ row)

### Presynaptic potentiation relies on the *cacophony* Ca_V_2.1 Ca^2+^ channel

Associative learning alters Ca^2+^ transients in MB γ neurons (Louis et al., 2018), which could influence neurotransmitter release. Major sources of stimulus-evoked intracellular Ca^2+^ include influx through voltage-sensitive Ca_V_2 channels, which are involved in presynaptic short-term and homeostatic plasticity (Frank et al., 2006; Inchauspe et al., 2004; Ishikawa et al., 2005; Muller and Davis, 2012). To probe the mechanisms of Ca^2+^-dependent molecular mechanisms underlying presynaptic plasticity, we first knocked down the α subunit of the Ca_V_2 Ca^2+^ channel encoded by *cacophony* (Cac), in the mushroom body. Cac was knocked down conditionally in adult MBs with RNAi using the R13F02-Gal4 driver, combined with the ubiquitous temperature-sensitive tub-Gal80^ts^ repressor (McGuire et al., 2003) to circumvent any potential for developmental effects (Fig. 3A). RNAi expression was induced four days prior to the experiment, and ACh release from MB neurons was imaged with GRAB-ACh (Jing et al., 2018; Zhang et al., 2019). Control flies (containing R13F02-Gal4, UAS-GRAB-ACh, and tub-Gal80^ts^, but lacking a UAS-RNAi) exhibited plasticity across the γ lobe in the same spatial patterns as previously observed: there was an increase in relative CS+ responses in the γ1-γ3 compartments, and a trend toward a CS+ decrease in γ5 (Fig. 3 C,E,F, S4). When Cac was knocked down conditionally, odor-evoked ACh release was still observed, demonstrating that synaptic exocytosis remained intact. Yet the CS+ potentiation was lost across the γ1 - γ3 compartments (Fig. 3 D,G, S4). This demonstrates that potentiation of ACh release to the trained odor – induced by learning – is dependent on the presynaptic Ca_V_2.1 channel Cac.

**Figure 3.**
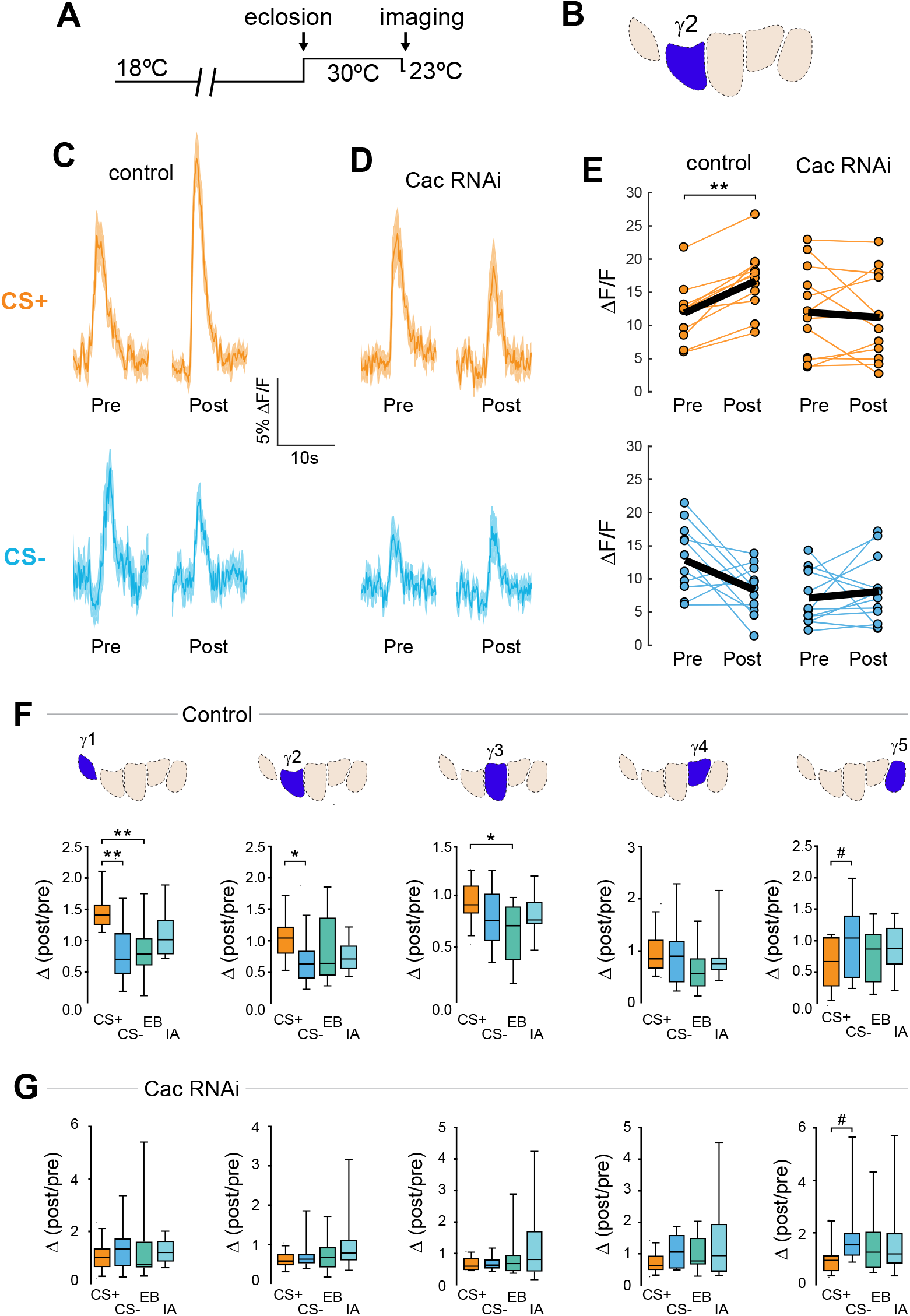
Conditional knockdown of the Ca_V_2 channel Cac impairs potentiation of ACh release from the MB following appetitive conditioning. **(A)** Diagram of the temperature shifts employed for conditional knockdown of Cac with tub-Gal80ts. **(B)** Diagram of the MB compartments, highlighting the γ2 compartment that was imaged for the data shown in panels C-E. **(C)** Pre- and post-conditioning CS+ (orange; top) and CS− (blue; bottom) odor-evoked ACh release from the γ1 compartment before and after appetitive conditioning, imaged in control animals (w;UAS-GRAB-ACh/+; R13F02-Gal4/UAS-tub-Gal80ts). Time series trace with line and shading representing mean ± S.E.M. **(D)** CS+ and CS− odor-evoked ACh release from the γ1 compartment in animals with conditional knockdown of Cac (w;UAS-GRAB-ACh/UAS-Cac-RNAi;R13F02-Gal4/UAS-tub-Gal80ts). **(E)** Pre- and post-conditioning ΔF/F CS+ and CS− responses in control and Cac knockdown animals. **(F)** Change in ACh release (post/pre response) following appetitive conditioning (CS+ and CS−) and odor-only presentation (EB: ethyl butyrate; IA: isoamyl acetate) in control animals across the five MB γ lobe compartments: γ1-γ5 (left to right). **(G)** Change in ACh release across the five MB compartments in animals with conditional knockdown of Cac. **p<0.01, *p<0.01; #p<0.07; n = 12.

Data from the appetitive conditioning experiments suggested that potentiation of the CS+ response was dependent on Cac. Interestingly, the trend toward CS+ depression in the most distal γ5 compartment remained intact when Cac was knocked down (Fig. 3). This suggests that presynaptic potentiation, but not depression, requires the voltage-sensitive Ca_V_2 Ca^2+^ channel *cacophony* across the MB compartments. To further examine whether depression of the CS+ was affected, we turned to aversive conditioning, which generates robust CS+ depression in the proximal γ compartments (Fig. 2). Control flies for conditional knockdown experiments exhibited similar CS+ depression in the proximal γ2,γ3 lobes. Knock down of Cac did not appreciably impair depression of CS+ responses. There was a significant depression in γ2, both in terms of CS+:CS− and CS+ relative to odor-only controls (Figs. 4, S5). In γ1, there was a trend toward a decrease in the CS+:CS− ratio that matched the controls (Fig. S5). In γ3, the difference between the CS+ and CS− (or odor-only control) did not reach significance, but there was a trend in the same direction as the controls (Fig. S5). Overall, these data demonstrate that Cac is not required for learning-induced depression of ACh release.

**Figure 4.**
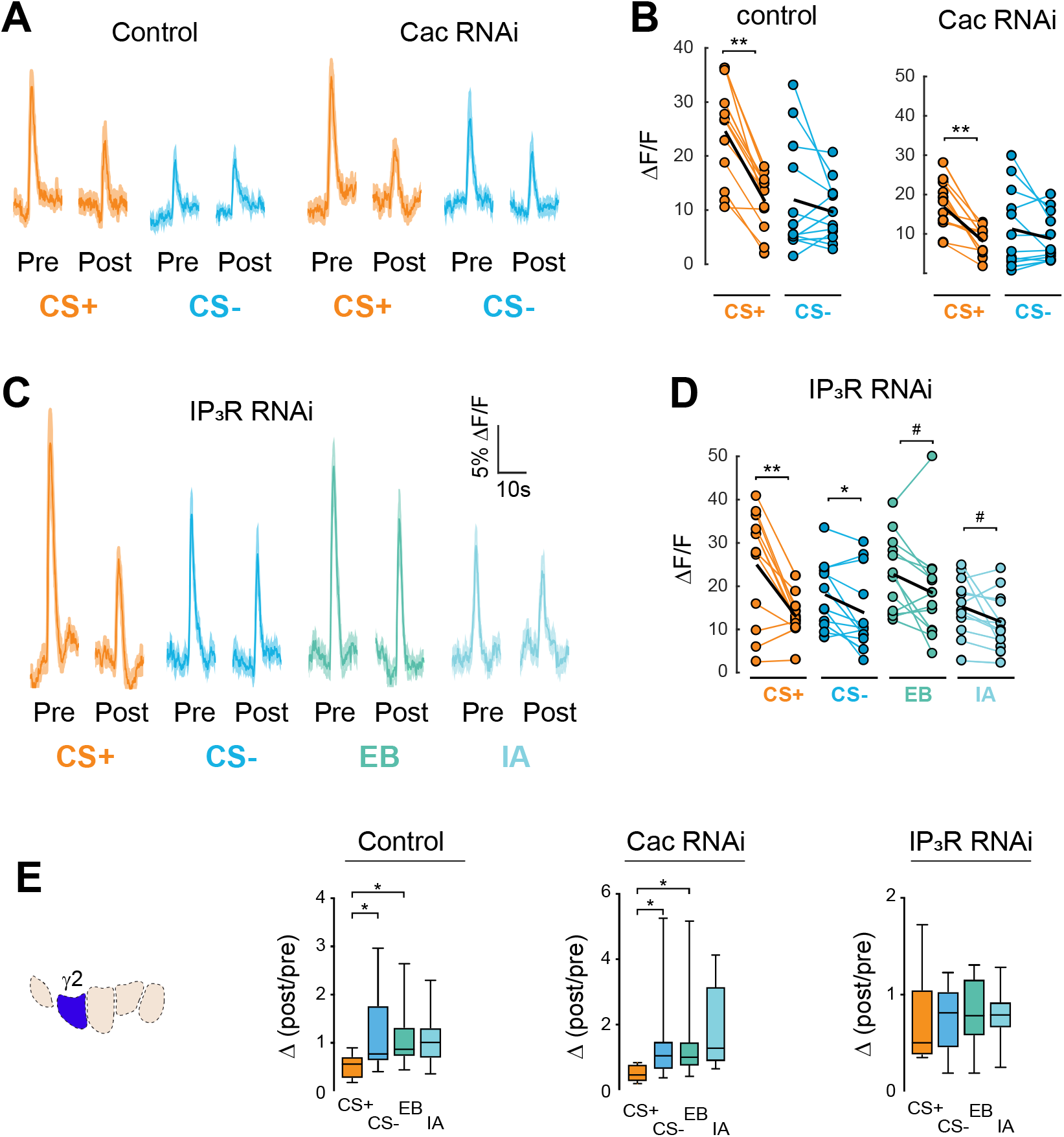
Cac and IP_3_R exert distinct effects on synaptic plasticity and maintenance of olfactory responses following aversive conditioning. **(A)** Pre- and post-conditioning CS+ and CS− odor-evoked ACh release in control and Cac RNAi flies. Time series trace with line and shading representing mean ± S.E.M. **(B)** Pre- and post-conditioning ΔF/F CS+ and CS− responses in control and Cac knockdown animals. Each thin line connects the pre- (left) and post-conditioning response (right) for one animal. The thick black line represents the mean. **(C)** ACh release in IP_3_R knockdown animals for trained odors (CS+ and CS−) as well as the respective odor-only controls (ethyl butyrate [EB] and isoamyl acetate [IA]). **(D)** ΔF/F responses in IP R knockdown flies. **(E)** Change in ACh release (post/pre response) following aversive conditioning (CS+ and CS−) and odor-only presentation (EB and IA) in controls, as well as flies with conditional Cac and IP_3_R knockdown.

### Post-conditioning odor contrast and maintenance of odor responses are dependent on IP_3_ signaling

Ca^2+^ release from the endoplasmic reticulum (ER) is a major source of stimulus-evoked Ca^2+^ in neurons, including MB neurons, and modulates various forms of synaptic/homeostatic plasticity (Handler et al., 2019; James et al., 2019; Taufiq et al., 2005). Therefore, we reasoned that inositol triphosphate receptor (IP_3_R) mediated Ca^2+^ release may contribute to presynaptic plasticity across MB compartments. To test this, we conditionally knocked down the IP_3_R in the adult MB with RNAi. GRAB-ACh was expressed in the MB (as above) while conditionally knocking down IP_3_R (Fig. 4, S5). For these experiments, flies were aversively conditioned (IP_3_R knockdown impairs feeding under the microscope, precluding appetitive conditioning).

Knockdown of IP_3_R eliminated the post-conditioning contrast between the CS+ and CS− (i.e., the difference between the CS+ and CS−) (Fig. 4, S5). This was due to increased adaptation to the odors (reduction in post-conditioning odor responses). This occurred in the CS− and both odor-only control groups, bringing them down to a similar level to the level of the CS+ group (Fig. 4 C-E). Thus, in normal conditions, release of Ca^2+^ from the ER via IP_3_R is necessary to maintain odor responsivity upon repeated odor presentations. Loss of IP_3_R renders the MB neurons more susceptible to adaptation, reducing the contrast between the CS+ – which exhibits depression following aversive learning – and the other odor(s).

### Compartmentalized plasticity propagates into downstream mushroom body output neurons

Since ACh release from each compartment provides input to unique postsynaptic mushroom body output neurons, the presynaptic plasticity observed in each compartment should be mirrored in the respective postsynaptic MBON(s) innervating that compartment. To test this, we imaged Ca^2+^ responses in MBONs with GCaMP and examined the effect of appetitive conditioning. Four sets MBONs were tested, each innervating and receiving cholinergic input from a distinct MB γ lobe compartment: γ1 pedc> α/β, γ2 α′1, γ3/γ3 β′1, and γ5β′2a (Fig. 5A).

**Figure 5.**
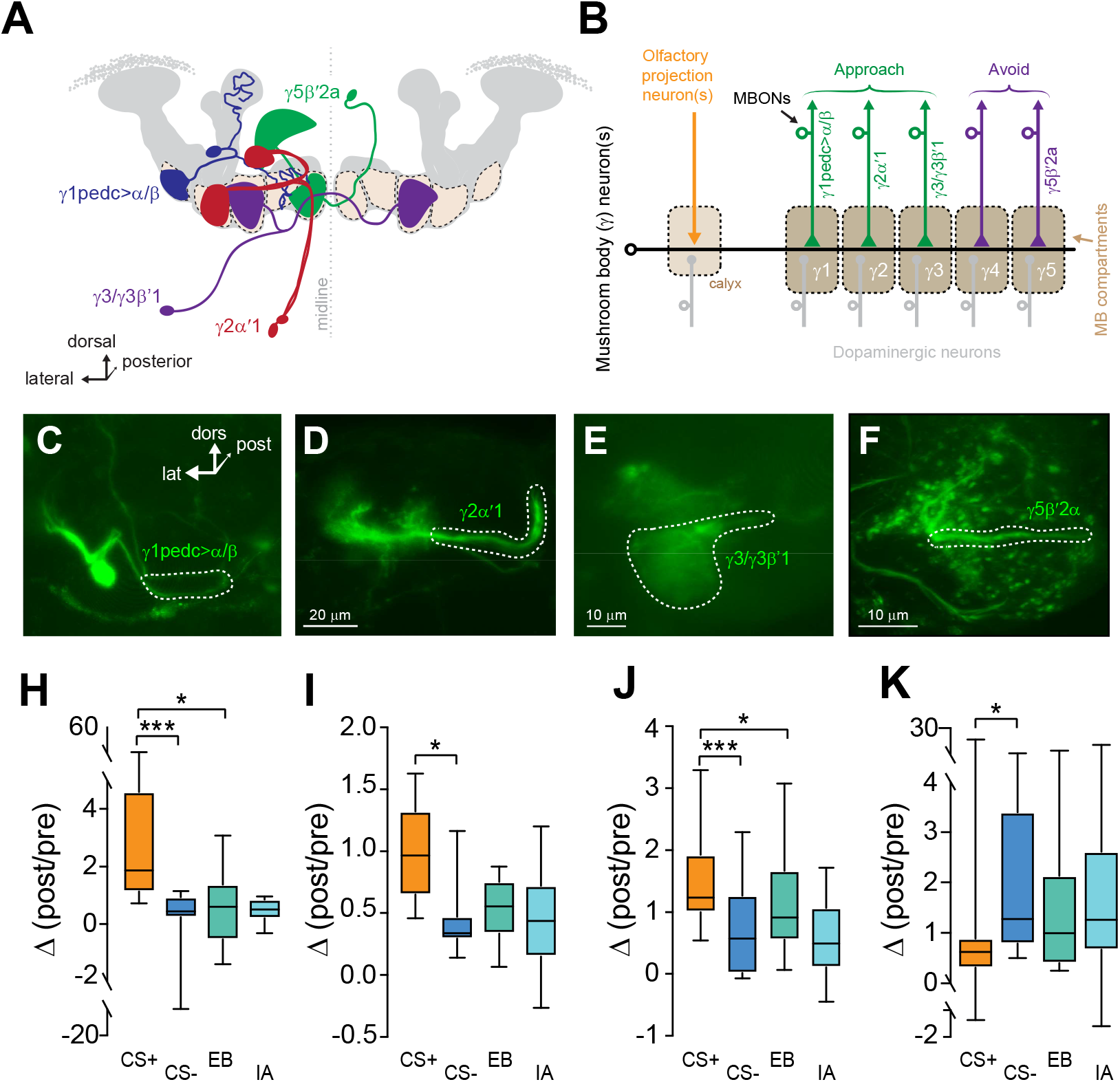
Plasticity in MBON Ca^2+^ responses mirrors compartmental plasticity in the MB neurons. **(A)** Diagram of MBONs innervating specific γ lobe compartments, viewed from a frontal plane. Each MBON is bilaterally paired, though only one is drawn here for visual clarity. **(B)** Circuit diagram of the dopaminergic neurons and MBONs in each compartment, as well as the putative valence associated with each compartment/MBON. **(C-F)** Diagrams of the γ1pedc> /β, γ2α′1, γ3, and γ5 β′2a MBONs, respectively, drawn unilaterally in isolation. **(G-J)** Representative confocal images of the γ1pedc> α/β, γ2 α′1, γ3, and γ5β ′2a MBONs, respectively. The region of interest circumscribed for quantification (neuropil or an efferent neurite) is drawn with a dotted white line. lat: lateral, dors: dorsal, post: posterior. **(K-N)** Change in odor-evoked responses (Post/pre responses), following conditioning (CS+ and CS−). ***p<0.001, *p<0.01; n = 12 (Kruskal-Wallis/Bonferroni).

Within the lobe, these neurons innervate the γ1, γ2, γ3, and γ5 compartments, respectively (Fig. 5 B-F). The γ1pedc> α/β MBON exhibited a significant elevation of the CS+ response relative to the CS− (↑CS+:CS−) (Fig. 5H). This was due to a potentiation of the CS+ response, as the post-conditioning CS+ response was significantly larger than the corresponding odor-only control. The γ2 α′1 MBON also exhibited an increase in the CS+:CS− ratio following conditioning (Fig. 5I). In this neuron, the plasticity could not be unambiguously attributed to purely CS+ potentiation or CS− depression. The γ3/γ3 β ′1 MBONs exhibited an increase in the CS+:CS− that was due to potentiation of the CS+ response (Fig. 5J). Note that these neurons are not parsed with available drivers and were imaged as a pair. Presynaptically, the γ3 compartment exhibited a depression in the CS− response, suggesting that the potentiation in the MBON CS+ response may emanate either from the β ‘1 inputs or modulation via polysynaptic circuit interactions.

Finally, appetitive conditioning produced plasticity in the opposite direction in the 5 β ′2a MBON; this neuron exhibited a decrease in the CS+ response relative to the CS− (↓CS+:CS−) (Fig. 5K). In each case, the directionality of the plasticity (CS+:CS−) matched that observed in ACh responses in the presynaptic compartment. Thus, compartmentalized, presynaptic plasticity in neurotransmitter release from the MB compartments likely plays a role in modulating the MBON responses following learning.

### Isolation of timing effects reveals CS− specific depression in the γ3 compartment

Synaptic depression in ACh release in the γ2 and γ3 compartments following appetitive conditioning was unique in that, in wild-type animals, it involved plasticity to the CS− (Figs. 2 G,H, S3). This raised the question of whether the simple act of presenting an odor 30 seconds after the offset of US pairing – the time at which the CS− is presented in the conditioning paradigm – is sufficient to alter ACh release. To test this, we compared the results from the discriminative CS+/CS− imaging assay (Figs. 2, S3) with a single-odor paradigm (Fig. 6A). Flies expressing GRAB-ACh in the MB via the 238Y-Gal4 driver were presented with an odor and sucrose, in a similar manner to the standard discriminative appetitive conditioning protocol, except that the CS+, CS−, or US was omitted (Fig. 6A, S6). We compared the change in responses to that odor across the three protocols in all five compartments (Fig. 6C, S3). This revealed several major facets of plasticity in ACh release following appetitive conditioning. First, discriminative training is necessary for the potentiation in γ1 and γ2, which was lost in single-odor CS/US training (protocol #1) (Fig. 6A,C, S6). In addition, when omitting the CS+, only the γ3 compartment revealed significant timing effects (Fig. 6 B,C, S6); presenting sucrose prior to presentation of an odor in the normal CS− time slot (protocol #2) resulted in a significantly smaller response than CS/US pairing, as well as a trend toward depression relative to the odor-only group. Therefore, the backward temporal contingency of the odor and sucrose presentation likely underlies the depression of odor-evoked responses in the γ3 region observed with discriminative CS+/CS− learning (Fig. 2H). Overall, these data demonstrate that the γ3 compartment is particularly important for the temporal comparison of the CS+ and CS−, which is critical for discriminative learning (a possibility we explore further below).

**Figure 6.**
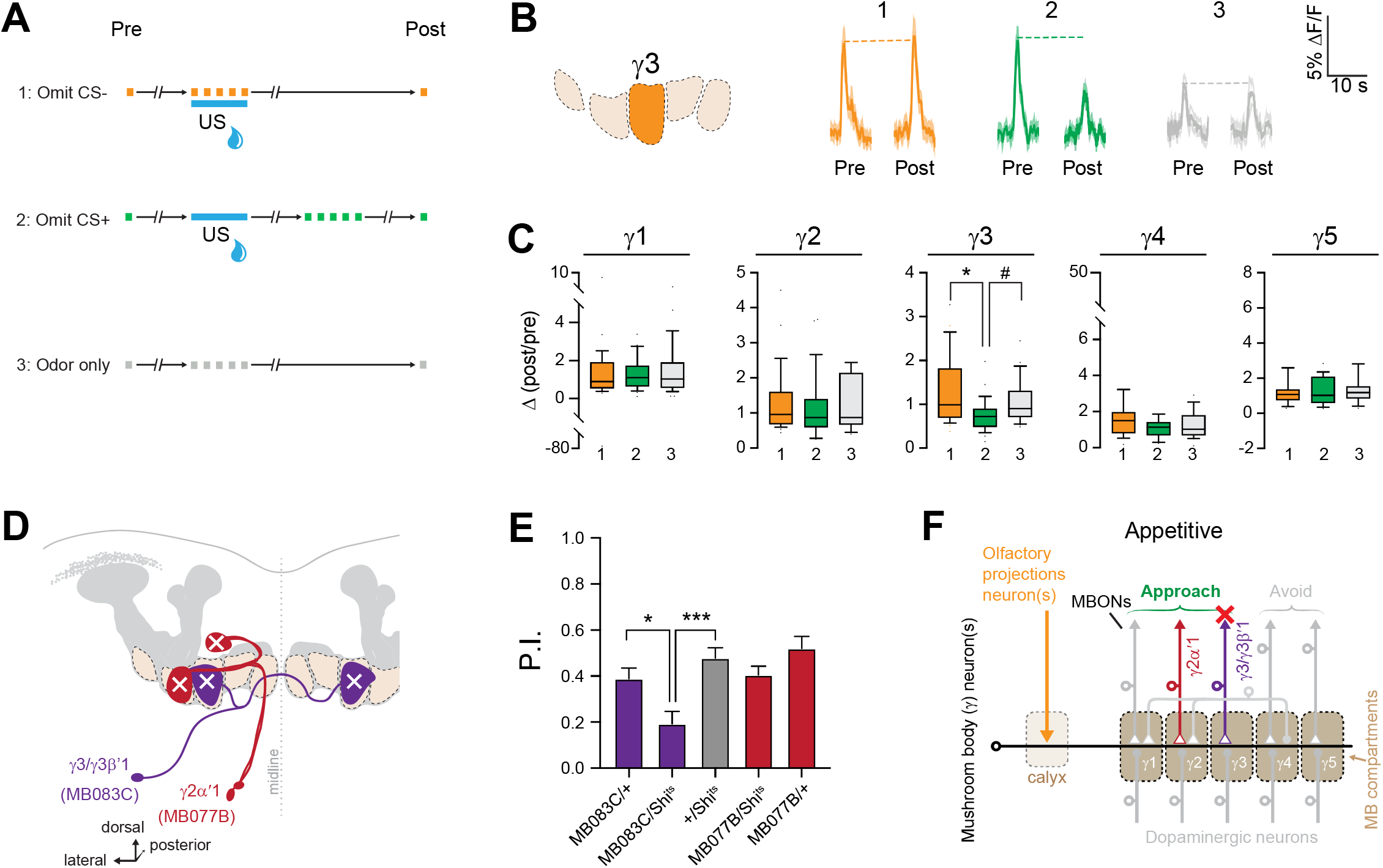
MB γ3 plasticity encodes appetitive timing (CS−) effects, and output neurons from this region are necessary for appetitive learning. **(A)** Diagram of the training paradigms utilized for ACh imaging experiments. **(B)** Time series traces showing odor-evoked GRAB-ACh responses from the γ3 compartment with all three protocols. **(C)** Quantification of the odor-evoked post/pre responses from each γ lobe compartment. *p<0.01, #p=0.016; n = 27 (Kruskal-Wallis/Bonferroni). **(D)** Anatomical Diagram of the γ2α′1 (MB077B-Gal4) and γ3/γ3 β′1 (MB083C-Gal4) MBONs. **(E)** Behavioral appetitive conditioning in flies, silencing either γ2 α′1 or γ3/γ3 β ′1 MBONs with Shibirets (Shits), compared to heterozygous Gal4/+ and UAS/+ controls. P.I.: Performance Index. *p < 0.05, ***p<0.0005 (ANOVA/Sidak); n = 16. **(F)** Circuit diagram of the output of the γ2 ′1 and γ3/γ3 β ′1 MBONs.

### Synaptic activity from the γ3-innervating MBONs mediate appetitive learning

The unique role of the γ2 and γ3 compartments in encoding CS− plasticity led us to question the behavioral roles of the MBONs that receive input from these compartments (Figs. 1, 6A-C). With the exception of the γ1pedc> α/β (Perisse et al., 2016), the involvement of these MBONs in appetitive memory is unclear. To test whether the MBONs innervating the γ2 and γ3 compartments mediate appetitive memory, we carried out behavioral appetitive classical conditioning, blocking synaptic transmission from MBONs with *Shibire*^ts^ (*Shi*^ts^) (McGuire et al., 2001) (Fig. 6 D,E). Blocking the γ2 α′1 MBON did not significantly impair performance in Appetitive conditioning. Therefore, while activation of the γ2 α′1 MBON drives approach behavior (Aso et al., 2014b) and the neuron is necessary for aversive memory (Berry et al., 2018), it is not crucial for appetitive learning in the otherwise intact nervous system. In contrast, blocking synaptic transmission from the γ3/γ3 β′1 MBONs significantly impaired appetitive conditioning performance (Fig. 6E). This demonstrates that the output of the γ3/γ3 β′1 MBONs is necessary for normal appetitive short-term memory (Fig. 6F). These neurons convey the output of the MB γ3 compartment to the crepine and superior medial protocerebrum (where they innervate interneurons that project to the fan-shaped body and lateral accessory lobe further downstream), as well as provide direct contralateral MB feedback and form polysynaptic feedback loops via MB-innervating PAM dopaminergic neurons and other MBONs (Scaplen et al., 2021; Xu et al., 2020). These multi-layered connections provide several routes through which they could modulate behavioral output following learning. Overall, the present data suggest that the γ3/γ3 β′1 MBONs receive input from an MB compartment with unique physiology, and represent a key node through which discriminative effects influence appetitive memory and decision-making.

## Discussion

Compartmentalized plasticity in neurotransmitter release expands the potential computational capacity of learning circuits. It allows a set of odor-coding mushroom body neurons to bifurcate their output to different downstream approach- and avoidance-driving downstream output neurons, independently modulating the synaptic connections to alter action selection based on the conditioned value of olfactory stimuli. The MB modifies the encoded value of olfactory stimuli through bidirectional plasticity in odor responses, which are compartment-specific along the axons. The CS+ and CS− drive unique patterns of plasticity in each compartment, demonstrating that olfactory stimuli are reweighted differently across compartments following learning, depending on the temporal associations of the stimuli. Different molecular mechanisms regulate the potentiation of trained odor responses (Ca_V_2/Cac) and maintenance of responsivity over time (IP_3_R). Finally, one set of γ output neurons, the γ3/γ3β ′1 MBONs, is particularly important for appetitive short-term memory.

The present data reveal learning-induced, bidirectional plasticity of ACh release in the MB neurons following conditioning with naturalistic stimuli *in vivo*, which was compartmentally-localized and coherent with the innate valence of the MBON innervating the compartment. Notably, the γ2 and γ3 compartments, which relay information to approach-promoting MBONs (Aso et al., 2014b), exhibited enhanced CS+:CS− responses to appetitive conditioning, and conversely reduced CS+:CS− following aversive conditioning. This was observed within 5 minutes of conditioning, a time point consistent with short-term memory in behavioral assays. Previous studies have described short-term, heterosynaptic depression in the γ1pedc MBON following reinforcement substitution via PPL1 dopaminergic neuron stimulation (Hige et al., 2015a) and changes in odor-evoked Ca^2+^ responses following olfactory classical conditioning (Perisse et al., 2016). Aversive conditioning has also been shown to decrease neurotransmitter release from the MB neurons (Zhang and Roman, 2013; Zhang et al., 2019). Indirect evidence, via Ca^2+^ imaging in presynaptic MB neurons, has suggested that increases in presynaptic neurotransmission could be associated with learning. Specifically, reinforcement substitution by pairing odor with stimulation of appetitive PAM dopaminergic neurons potentiates odor-evoked cytosolic Ca^2+^ transients across the MB (Boto et al., 2014). In addition, appetitive conditioning with naturalistic odor + sucrose pairing increases odor-evoked cytosolic Ca^2+^ transients in MB neurons (Louis et al., 2018). However, all MB compartments exhibit plasticity with uniform directionality; short-term aversive conditioning produces no detectable change and appetitive conditioning uniformly elevates odor-evoked responses across the γ1-γ5 compartments. Thus, this effect is not selective for subcellular MB compartments that connect to the “aversive” or “appetitive” MBONs. More compartmentalized effects have been observed with other manipulations – presentation of sucrose alters synaptically-localized Ca^2+^ transients in a compartmentalized manner (Cohn et al., 2015), as does stimulation of γ4-innervating dopaminergic circuits (Handler et al., 2019).

This study revealed two major mechanisms regulating the spatial patterns of compartmentalized plasticity across the MB compartments: a Cac-dependent CS+ potentiation and an IP_3_R-dependent maintenance of sensory responses. This suggests that different sources of intracellular Ca^2+^ play different roles in regulating MB synaptic responses. Cac is the pore-forming subunit of the voltage-sensitive, presynaptic Ca_V_2 Ca^2+^ channel in *Drosophila*. Ca_V_2 channels regulate several forms of synaptic plasticity, including paired-pulse facilitation, homeostatic plasticity, and long-term potentiation (Frank et al., 2006; Inchauspe et al., 2004; Nanou et al., 2016). Our data suggests that these channels regulate the spatial patterns of learning-induced plasticity in the MB unidirectionally, with Cac underlying potentiation but not depression. Ca_V_2 channel activity is modulated by presynaptic calcium and G protein-coupled receptor activity (Zamponi and Currie, 2013), and channel localization in the active zone dynamically regulates synaptic strength (Gratz et al., 2019; Lubbert et al., 2019). These mechanisms may play a role in increasing CS+ responses following appetitive conditioning, as activity in MB neurons results in increased intracellular Ca^2+^ and dopaminergic neurons innervating the MB activate receptors that are important for memory formation (Boto et al., 2014; Cohn et al., 2015; Gervasi et al., 2010; Kim et al., 2007; Schwaerzel et al., 2003; Tomchik and Davis, 2009). Baseline stimulus-evoked neurotransmitter release in Cac knockdown was maintained, likely mediated by either residual Cac expression or compensation by other intracellular Ca^2+^ channels/sources. In contrast to potentiation, IP_3_R was necessary to maintain normal odor responsivity when odors were presented on multiple trials (i.e., across pre/post odor presentations). This is consistent with the role of IP_3_R in maintenance of presynaptic homeostatic potentiation at the neuromuscular junction (James et al., 2019). In addition, dopaminergic circuits associated with reward learning drive release of Ca^2+^ from the endoplasmic reticulum when activated with MB neurons in a backward pairing paradigm *ex vivo*, potentiating MB γ4 connections with the respective γ0034 MBON (Handler et al., 2019).

Alterations of MBON activity following learning are likely the product of both synaptic plasticity at the MB-MBON synapses and indirect circuit effects, such as feedforward inhibition (Aso et al., 2014a; Cervantes-Sandoval et al., 2017; Perisse et al., 2016). Polysynaptic inhibitory interactions can convert depression from select MB compartments into potentiation in MBONs following learning. In one established example, reduction of odor-evoked responses in the GABAergic γ1pedc MBON following aversive conditioning disinhibits the downstream γ5 ′2a MBON (Owald et al., 2015; Perisse et al., 2016). It is unclear whether this mechanism generalizes to other MB compartments. The present data demonstrates that learning drives potentiation and depression of ACh release across multiple MB compartments, providing a direct mechanism for altering MBON responses. Importantly, by comparing the CS+ and CS− responses to those of untrained odors, we ascribed differences between the CS+ and CS− to potentiation or depression in absolute terms within each compartment. This uncovered an additional layer of spatial regulation of plasticity in the γ1-γ3 compartments: a gradient of CS+ potentiation to CS− depression following appetitive conditioning, which is elaborated in greater detail below. In addition, it revealed that the IP_3_-dependent loss of CS+/CS− contrast was due, at least in large part, to alterations in olfactory adaptation.

The CS+/CS− relationship changed in a linear gradient down the γ1-γ3 compartments following appetitive conditioning. Appetitive conditioning increased CS+ responses in the γ1 compartment, while decreasing the CS− responses in the γ3 compartment. The γ2 compartment yielded a mix of these responses. These patterns of plasticity have the net effect of increasing the relative response to the CS+ odor. Since the MBONs postsynaptic to these compartments drive behavioral approach (Aso et al., 2014b), these patterns of plasticity would bias the animal’s behavior toward approach of the CS+ if the animal faced both odors simultaneously.

Such a situation would occur at the choice point of a T-maze during retrieval in a classical conditioning assay. This further suggests loci where for CS+ and CS− plasticity, which are suggested by behavioral data indicating that temporal/CS− information contribute to behavioral memory (Handler et al., 2019; Konig et al., 2018; Tanimoto et al., 2004; Tully and Quinn, 1985). This is physiologically reflected in plasticity in ACh release to the CS+ and/or CS− across multiple compartments. For instance, the γ2 and γ3 compartments exhibited a depression in ACh release to the CS−. Therefore, consequences to the specific timing of odor-evoked responses prior to or after the delivery of the US play a key role in memory formation, with bidirectional plasticity forming within the MB neurons based on timing events, valence of the US, and local dopamine signaling (Handler et al., 2019; Konig et al., 2018; Tanimoto et al., 2004; Yamagata et al., 2016).

MBONs innervating the γ lobe drive approach/avoidance behavior when stimulated (Aso et al., 2014b). Despite the approach-promoting valence of the γ2 ′1 and γ3/γ3 β′1 MBONs, only the γ3/γ3 β′1 produced a loss-of-function phenotype in appetitive conditioning. This suggests that either the γ2 α′1 MBONs are uninvolved in appetitive learning (despite exhibiting learning-related plasticity), or that redundancy and/or different weighting across approach-promoting MBONs, renders the system resilient to silencing some of them. Blocking synaptic output of γ3/γ3 β′1 reduced appetitive conditioning performance, suggesting that these neurons play a particularly important role in appetitive learning.

Overall, plasticity between MB neurons and MBONs may guide behavior through biasing network activation to alter action selection in a probabilistic manner. Appetitive conditioning drives compartmentalized, presynaptic plasticity in MB neurons that correlates with postsynaptic changes in MBONs that guide learned behaviors. Prior studies documented only depression at these synapses at short time points following conditioning (Hige et al., 2015a; Zhang and Roman, 2013; Zhang et al., 2019). Here we observed both potentiation and depression in ACh release in the MB, suggesting that bidirectional presynaptic plasticity modulates learned behaviors. These bidirectional changes likely integrate with plasticity at downstream circuit nodes that also undergo learning-induced plasticity to produce network-level alterations in odor responses across the olfactory pathway following salient events. Thus, plasticity in ACh release from MB neurons function to modulate responsivity to olfactory stimuli features across graded plasticity maps down the mushroom body axons.

## Materials and Methods

### Fly Strains

Flies were fed and maintained on a standard cornmeal agar food mixture on a 12:12 light:dark cycle. The 238Y-Gal4 driver was selected for expression intensity in MB neurons (Louis et al., 2018). MBON drivers were selected from the FlyLight and split-Gal4 collections (R12G04, MB077b, and MB083c) (Jenett et al., 2012; Pfeiffer et al., 2010). The γ5 β′2a LexA MBON driver was a generated by Krystyna Keleman (Zhao et al., 2018). RNAi lines were obtained from the VDRC (Cac: 101478) (Dietzl et al., 2007) and TRiP collections (IP_3_R/*itpr*: 25937) (Perkins et al., 2015) and crossed into flies expressing R13F02-Gal4 and tub-Gal80^ts^ (McGuire et al., 2003). Final experimental genotypes were: Cac (w;UAS-GRAB-ACh/UAS-Cac-RNAi;R13F02-Gal4/UAS-tub-Gal80^ts^) and IP_3_R (w, UAS-GRAB-ACh/UAS-tub-Gal80^ts^;R13F02-Gal4;UAS-IP3R-RNAi), compared to genetic controls (w; UAS-GRAB-ACh/+;R13F02-Gal4/UAS-tub-Gal80^ts^).

### Fly preparation for *in vivo* Ca^2+^ imaging

Flies were briefly anesthetized, placed in a polycarbonate imaging chamber, and fixed with myristic acid (Sigma-Aldrich). The proboscis was fixed in the retracted position, except for appetitive conditioning experiments (as noted below). A cuticle window was opened, and the fat and tracheal air sacs were carefully removed to allow optical access to the brain. The top of the chamber was filled with saline solution (103 mM NaCl, 3mM MBl, 5mM HEPES, 1.5 mM CaCl_2_, 4 mM MgCl2·6H_2_O, 26 mM NaHCO_3_, 1 mM NaH_2_PO_4_·H2O, 10 mM trehalose, 7 mM sucrose, and 10 mM glucose), which was perfused over the dorsal head/brain at 2 mL/min via a peristaltic pump.

### *In vivo* imaging

GRAB-ACh (Jing et al., 2019; Jing et al., 2018; Zhang et al., 2019) was driven in the MB neurons, using the 238Y driver. Within the MB neurons, ROIs were drawn around five γ lobe compartments (γ1-5) within a single imaging plane for appetitive, and (γ2-5) for aversive. Imaging was performed with a Leica TCS SP8 confocal microscope utilizing appropriate laser lines and emission filter settings. Odors were delivered with an airstream for 1s (60mL/min flow rate) by directing the air flow with solenoid valves between an empty vial (air) to another containing 1μL odorant spotted on filter paper. Odor-evoked responses were calculated as the baseline normalized change in fluorescence (ΔF/F), using the maximum ΔF/F within a 4-s after odor delivery. The ratio of the post/pre responses were calculated as the maximum ΔF/F in an 8-s response window after odor delivery. In experiments with RNAi, flies expressing GRAB-ACh, a UAS-RNAi line, and tub-Gal80^ts^ were constructed; flies were raised at 18°C until eclosion, flies were transferred to 32°C 4-10 days prior to the experiment.

Experiments were carried out at room temperature (23°C) for ACh imaging/conditioning. For Ca^2+^ imaging experiments, GCaMP6f was expressed in the MBONs using the R12G04 (γ1pedc), MB077b (γ2 α ′1), MB083c (γ3) and VT014702 (γ5 β′2) Gal4 drivers. Experiments were carried out same as ACh imaging, except presenting a 3s odor delivery.

### Appetitive conditioning and imaging

Appetitive conditioning was carried out as previously described (Louis et al., 2018). One odor (the CS+) was presented in conjunction with a paired sucrose (1M, containing green food coloring) unconditioned stimulus (US), and a second odor (the CS−) was presented 30-s later. Both the CS+ and CS− odors were presented during conditioning experiments. In odor-only control cohorts, the sucrose US was omitted. During training, each odor (and the US) was presented continuously for 30 s for Ca^2+^ imaging experiments. Six 1-s odor pulses were presented during conditioning over a 30-s period, with a 5-s inter-pulse interval, to prevent desensitization of the reporter. Pre/post odor-evoked responses were imaged prior to and after the imaging protocol, using a 3-s (Ca^2+^ imaging) or 1-s (ACh imaging) odor pulse. During odor-evoked response imaging, proboscis extension was blocked utilizing a thin metal loop attached to a custom motorized micromanipulator. Flies were starved for a period of 18-24 hrs prior to conditioning. During conditioning, the proboscis was released, and the flies were presented sucrose through a metal pipette fed by a syringe pump controlled via a micro-controller (Arduino). To assess feeding, flies were monitored using a digital microscope (Vividia); sucrose ingestion was visually confirmed by the presence of green food coloring in the abdomen.

### Aversive conditioning and imaging

Flies were mounted in an aversive conditioning chamber such that the brain could be imaged while odors were delivered to the antennae and electric shocks delivered to the legs via a shock grid below the fly. Conditioning was carried out by pairing a CS+ odor with electric shocks as follows: 6x 1-s odor pulses, with a 5-s inter-pulse interval, paired with 6x 90-V electric shocks, followed 30s later by presentation of 6x 1-s pulses of the CS− odor with 5s inter-pulse interval. Pre- and post-conditioning odor-evoked responses were imaged using a 1-s odor pulse. In each animal, either the CS+ or CS− odor was tested pre- and post-conditioning.

### Behavioral appetitive conditioning

Adult flies, 2-5 day old, were trained under dim red light at 75% relative humidity. Appetitive conditioning experiments were performed in animals starved 16-20 h. Groups of ∼60 flies were exposed for 2 min to an odor (the CS−), followed by 30 s of air and 2 min of another odor, the (the CS+), paired with a 1M sucrose solution dried on filter paper, at 32°C for *Shibire*^ts^ blockade. The odor pairs were ethyl butyrate and isoamyl acetate, adjusted so that naive flies equally avoided the two odors (0.05 – 0.1%). Memory was tested by inserting the trained flies into a T-maze, in which they chose between an arm containing the CS+ odor and an arm containing the CS− odor. Flies were allowed to distribute for a 2 min choice period. The Performance Index (P.I.), calculated as (flies in the CS− arm)-(flies in the CS+ arm)/(total flies in both arms).

### Immunohistochemistry

5-7 days old adult flies were dissected in 1% paraformaldehyde in S2 medium, and processed according to a published protocol (Jenett et al., 2012). Brains and were incubated with the primary antibodies for 3 hours at room temperature and with the secondary antibodies for 4 days at 4°C. Incubations were performed in blocking serum (3% normal goat serum). Labeled brains were mounted in Vectashield media. Antibodies used were rabbit anti-GFP (1:1000, Invitrogen), mouse anti-brp (nc82) (1:50, DSHB), mouse anti-neuroglian (1:50,DSHB), goat anti-rabbit IgG and goat anti-mouse IgG (1:800, Alexa 488 or Alexa 633 respectively, Invitrogen). Images were obtained using Leica TCS SP8 confocal microscope.

### Quantification and Statistical Analysis

Data were compared with ANOVA/Sidak (parametric) or Kruskal-Wallis/Bonferroni (nonparametric) tests. Box plots show graph the median as a line, the 1^st^ and 3^rd^ quartile enclosed in the box, and whiskers extending from the 10^th^ to the 90^th^ percentile.

## Acknowledgments

The authors thank Krystyna Keleman for fly stocks, and Brock Grill for helpful discussions. Stocks obtained from the Bloomington Drosophila Stock Center (NIH P40OD018537) were used in this study. We thank Yuexuan Li for the help in the development of the GRAB-ACh sensor.

Research support was provided by NIH R00MH092294, R01 NS097237, the Whitehall Foundation (S.M.T.), NIH R35NS097224 (R.L.D.), the Beijing Municipal Science & Technology Commission Z181100001318002 (Y.L.), the Beijing Brain Initiative of Beijing Municipal Science & Technology Commission Z181100001518004 (Y.L.), Guangdong Grant “Key Technologies for Treatment of Brain Disorders” 2018B030332001 (Y.L.), the General Program of National Natural Science Foundation of China projects 31671118, 31871087, and 31925017) (Y.L.), the NIH BRAIN Initiative NS103558 (Y.L.), grants from the Peking-Tsinghua Center for Life Sciences (Y.L.) and the State Key Laboratory of Membrane Biology at Peking University School of Life Sciences (Y. L.).

## Competing Interests

The authors declare no competing financial interests.

---

**Figure S1.**
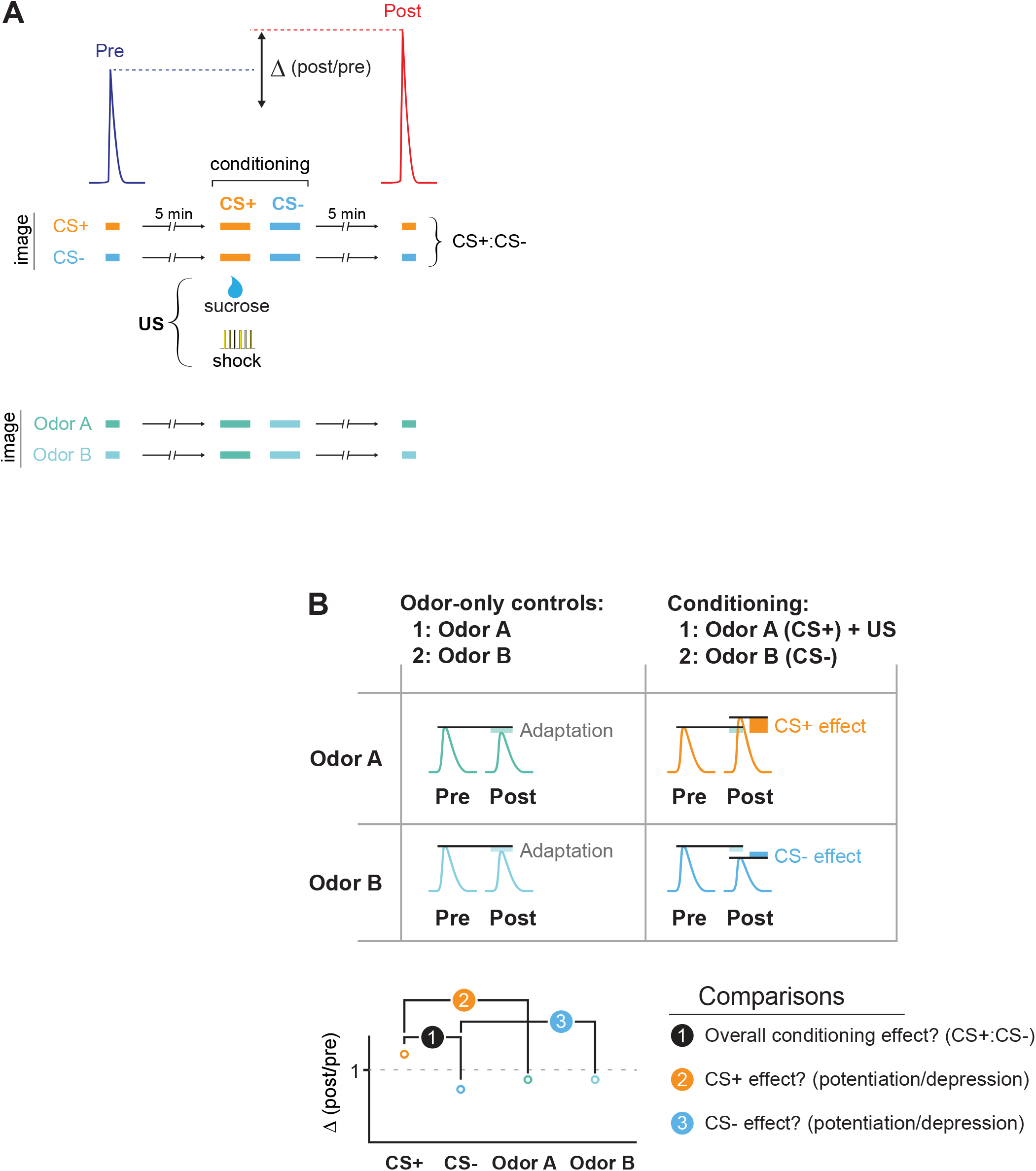
related to Figure 1. The conditioning and imaging assay and data analysis. **(A)** Flies were conditioned by pairing an odor (the CS+) with a US (electric shock or sucrose reward), and a second odor (the CS−) was presented afterward. Odor-evoked GACh or GCaMP responses were imaged in the MB and compared before (Pre) and after (Post) conditioning. Responses were compared to animals in which the same odors were presented, but no US presented (odor-only controls). To examine how conditioning (or odor-only presentation) changed the odor responses, the Δ (post/pre) was calculated for each treatment. **(B)** Two types of comparisons were made across conditions. First, we analyzed the CS+:CS− ratio, which mimics the putative comparison the animal makes when comparing the two odors at the choice point in a T-maze during memory retrieval. Second, we compared the CS+ and CS− to their respective odor-only controls in order to determine whether the responses were potentiated or depressed by conditioning. This comparison normalizes for any olfactory adaptation that is induced by the odor presentation during the training window.

**Figure S2.**
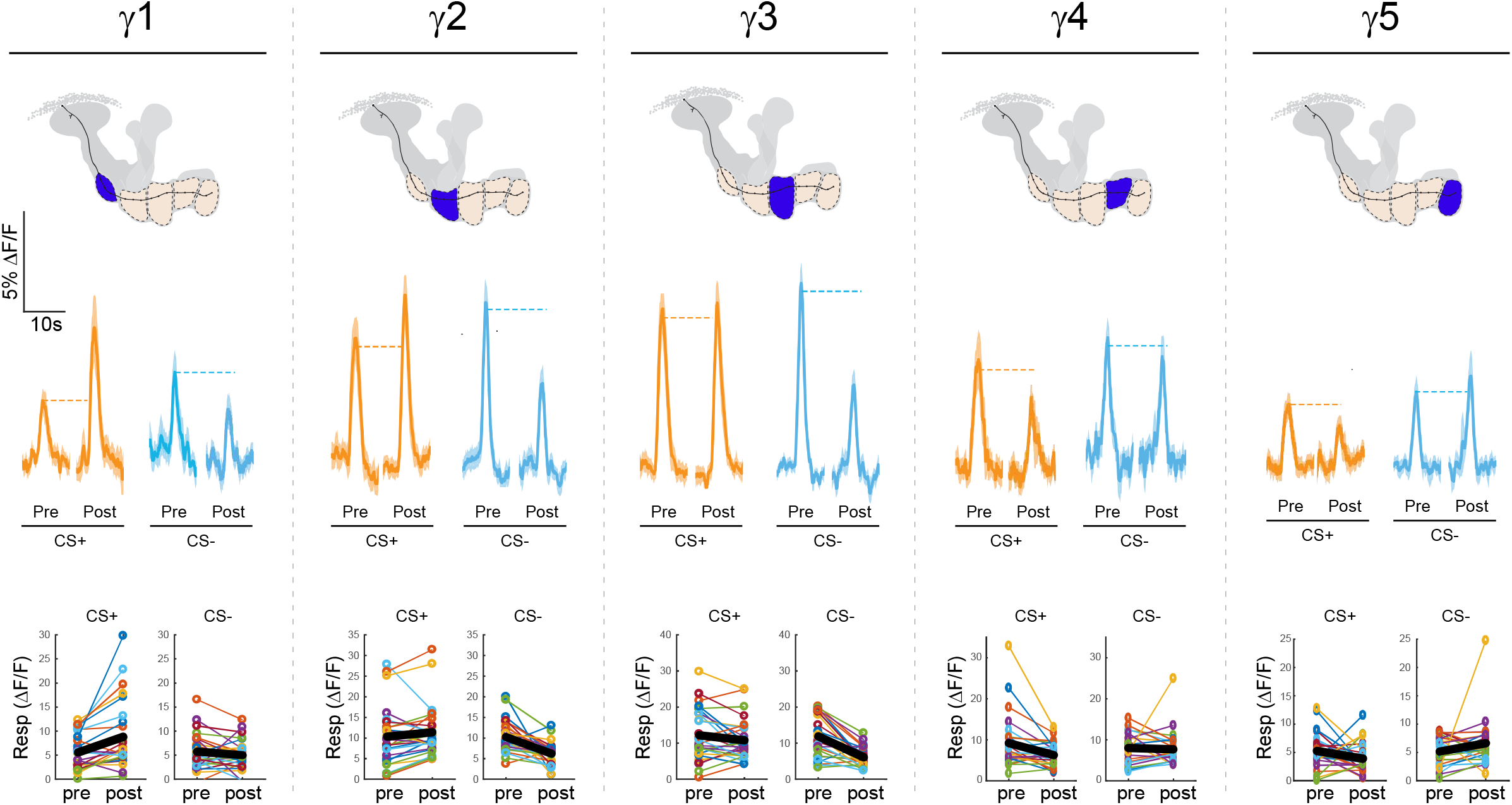
related to Figure 1. Effects of appetitive conditioning on GRAB-ACh responses across the γ lobe compartments. The top row shows diagrams of the location of each compartment within the mushroom body. The second row shows time series traces pre- and post-conditioning for the CS+ (ethyl butyrate [EB]) and CS− (isoamyl acetate [IA]). The third row shows quantification of the peak pre- and post-conditioning responses for each animal (n = 27). The thick black line represents the mean.

**Figure S3.**
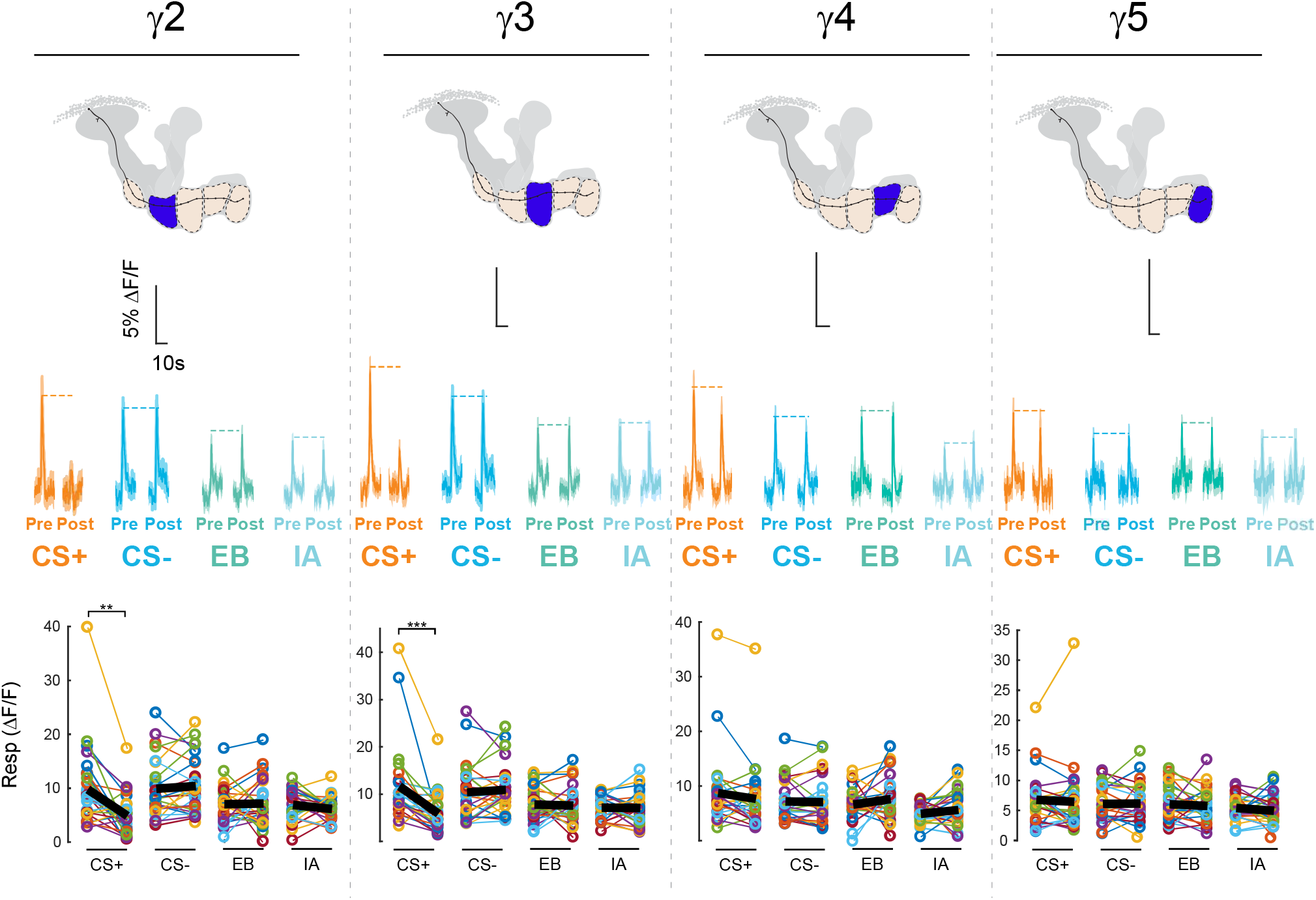
related to Figure 2. Effects of averisve conditioning on GRAB-ACh responses across the γ lobe compartments. The top row shows diagrams of the location of each compartment within the mushroom body. The second row shows time series traces pre- and post-conditioning for the CS+ (ethyl butyrate [EB]) and CS− (isoamyl acetate [IA]) and odor only controls. The third row shows quantification of the peak pre- and post-conditioning responses for each animal (n = 27).*p<0.05,*p<0.005, ***p<0.0005 n=12 (Wilcoxon rank-sum test) The thick black line represents the mean.

**Figure S4.**
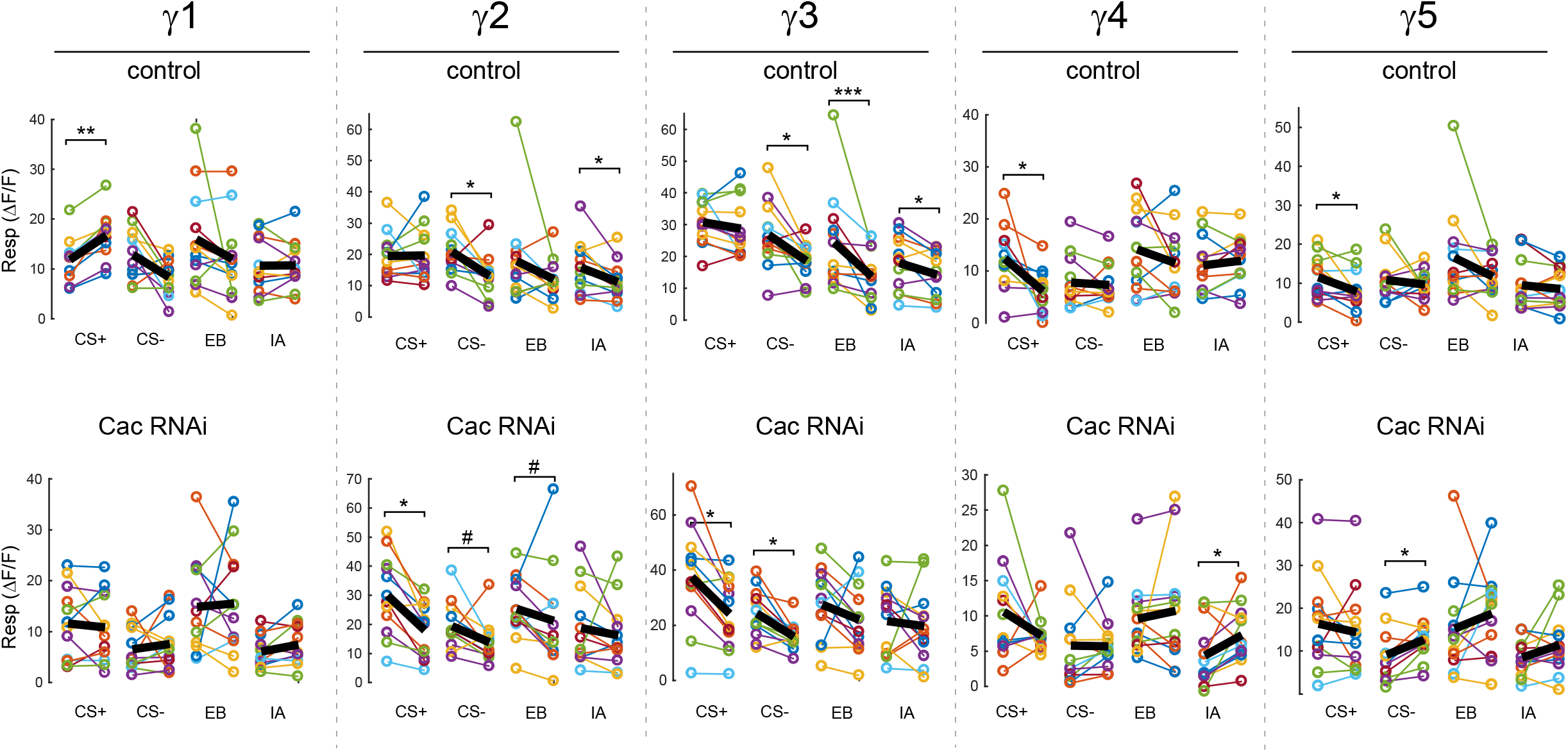
related to Figure 3. Effects of appetitive conditioning on GRAB-ACh responses across the γ lobe compartments using GRAB-ACh with control and cacophony knockdown flies. The top row shows time series traces pre- and post-conditioning for the CS+ (ethyl butyrate [EB]) and CS− (isoamyl acetate [IA]) of control flies. The quantification of the peak pre- and post-conditioning responses for each animal (n = 12) of control flies. *p<0.05,*p<0.005, ***p<0.0005, #p<0.07, n=12 (Wilcoxon rank-sum test). The second row shows time series traces pre- and post-conditioning Cac knockdowns.

**Figure S5.**
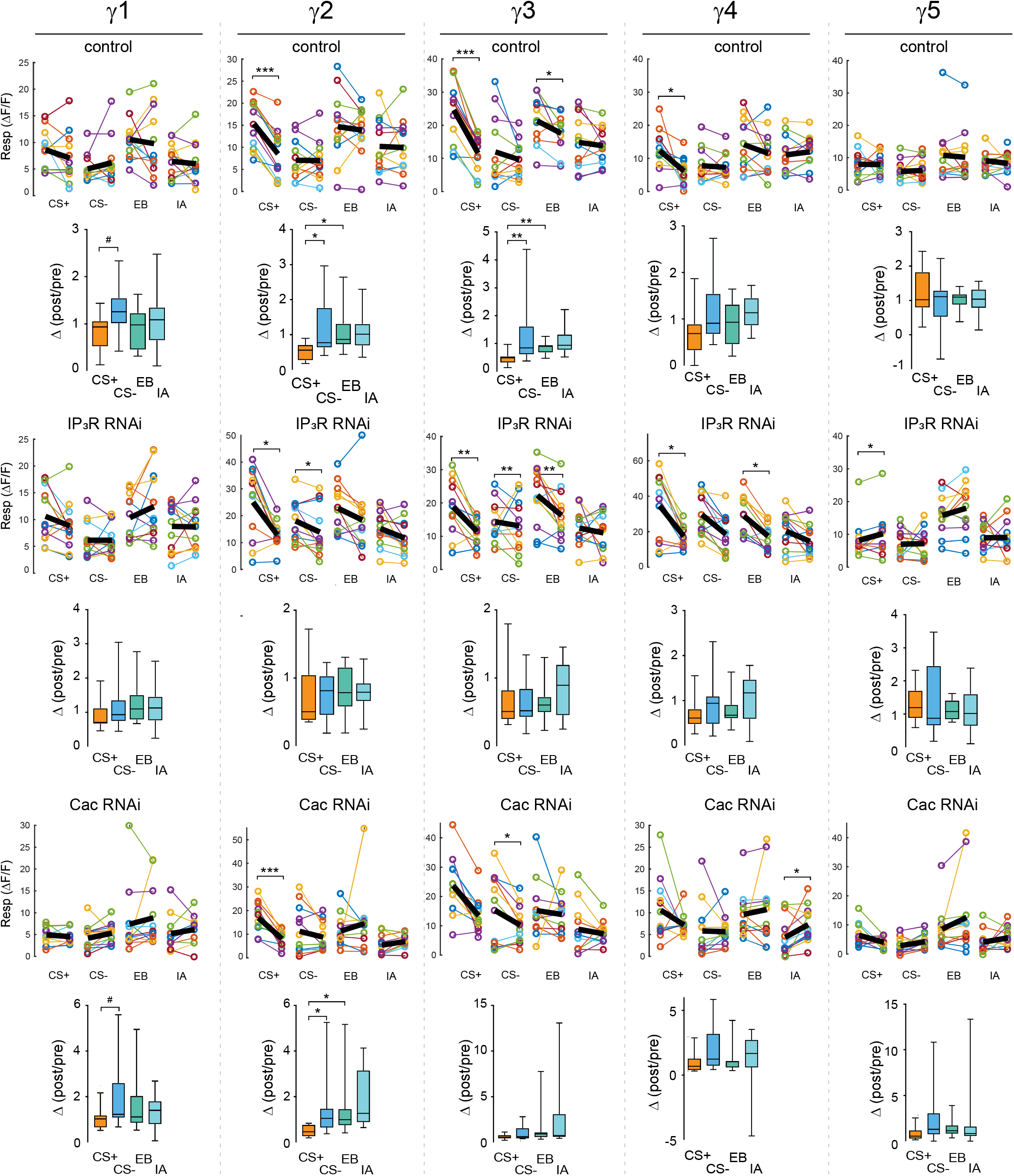
related to Figure 4. Effects of aversive conditioning on GRAB-ACh responses across the γ lobe compartments using GRAB-ACh with control, cacophony and IP_3_R knockdown flies. For all genotypes sample sizes, n=12 with statistical analysis (Wilcoxon rank-sum test) *p<0.05,*p<0.005, ***p<0.0005 for time series traces. For comparisons of CS+, CS−, and odor-only control responses (Kruskal-Wallis/Bonferroni) #p<0.03, *p<0.01, **p<0.001, ***p<0.0001. The top row shows time series traces pre- and post-conditioning for the CS+ (ethyl butyrate [EB]) and CS− (isoamyl acetate [IA]) and odor only, and the thick black line represents the mean. The second row shows comparisons of the CS+, CS−, and odor-only controls. The third row shows time series traces pre-post conditioning for IP_3_R knockdowns. The fourth row shows comparisons between the four treatments of IP3R knockdowns. The fifth row shows time series traces pre-post conditioning for Cac knockdowns. The final row shows comparisons between the four treatments of Cac knockdowns.

**Figure S6.**
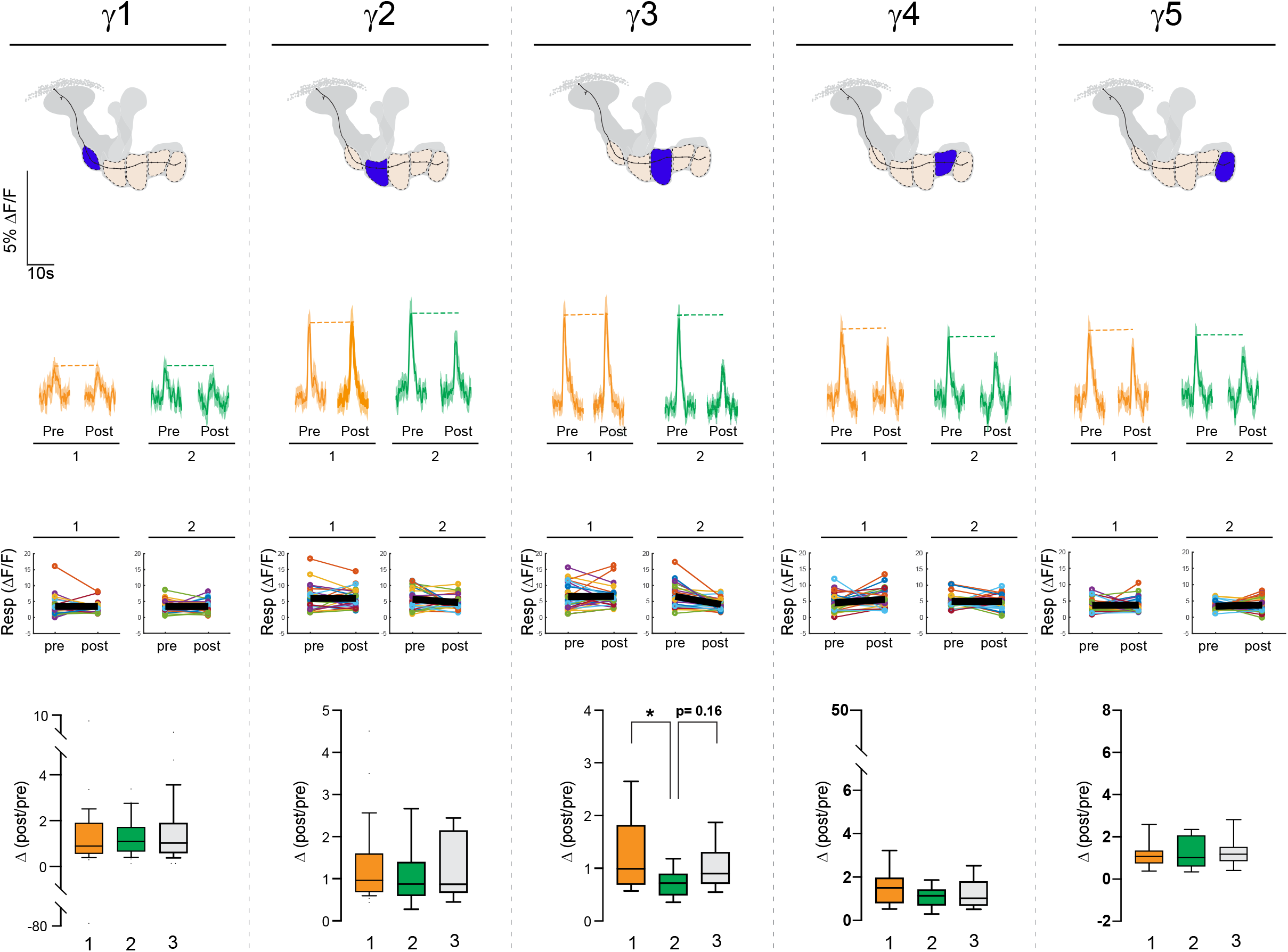
related to Figure 6. Effects of appetitive conditioning on GRAB-ACh responses across the γ lobe in the absence of either CS+ (1) or CS− (2). The top row shows diagrams of the location of each compartment within the mushroom body. The second row shows time series traces pre- and post-conditioning for paired, unpaired, and odor-only conditioning. The third row shows quantification of the peak pre- and post-conditioning responses for each animal (n = 27). The thick black line represents the mean. The bottom row shows comparisons of the CS+, CS−, and odor-only controls (EB and IA). *p<0.01, **p<0.001, ***p<0.0001; n =27 (Kruskal-Wallis/Bonferroni).

## Notes

### Competing Interest Statement

The authors have declared no competing interest.

## References

Aso, Y., Hattori, D., Yu, Y., Johnston, R.M., Iyer, N.A., Ngo, T.T., Dionne, H., Abbott, L., Axel, R., Tanimoto, H., and Rubin, G.M. (2014a). The neuronal architecture of the mushroom body provides a logic for associative learning. Elife 3.

Aso, Y., Sitaraman, D., Ichinose, T., Kaun, K.R., Vogt, K., Belliart-Guerin, G., Placais, P.Y., Robie, A.A., Yamagata, N., Schnaitmann, C., et al. (2014b). Mushroom body output neurons encode valence and guide memory-based action selection in Drosophila. Elife 3.

Barnstedt, O., Owald, D., Felsenberg, J., Brain, R., Moszynski, J.P., Talbot, C.B., Perrat, P.N., and Waddell, S. (2016). Memory-Relevant Mushroom Body Output Synapses Are Cholinergic. Neuron 89, 1237–1247.

Berry, J.A., Phan, A., and Davis, R.L. (2018). Dopamine Neurons Mediate Learning and Forgetting through Bidirectional Modulation of a Memory Trace. Cell Rep 25, 651–662 e655.

Boto, T., Louis, T., Jindachomthong, K., Jalink, K., and Tomchik, S.M. (2014). Dopaminergic Modulation of cAMP Drives Nonlinear Plasticity across the Drosophila Mushroom Body Lobes. Curr Biol 24, 822–831.

Boto, T., Stahl, A., Zhang, X., Louis, T., and Tomchik, S.M. (2019). Independent Contributions of Discrete Dopaminergic Circuits to Cellular Plasticity, Memory Strength, and Valence in Drosophila. Cell Rep 27, 2014–2021 e2012.

Cervantes-Sandoval, I., Phan, A., Chakraborty, M., and Davis, R.L. (2017). Reciprocal synapses between mushroom body and dopamine neurons form a positive feedback loop required for learning. Elife 6.

Cohn, R., Morantte, I., and Ruta, V. (2015). Coordinated and Compartmentalized Neuromodulation Shapes Sensory Processing in Drosophila. Cell 163, 1742–1755.

Crittenden, J.R., Skoulakis, E.M., Han, K.A., Kalderon, D., and Davis, R.L. (1998). Tripartite mushroom body architecture revealed by antigenic markers. Learn Mem 5, 38–51.

Dietzl, G., Chen, D., Schnorrer, F., Su, K.C., Barinova, Y., Fellner, M., Gasser, B., Kinsey, K., Oppel, S., Scheiblauer, S., et al. (2007). A genome-wide transgenic RNAi library for conditional gene inactivation in Drosophila. Nature 448, 151–156.

Frank, C.A., Kennedy, M.J., Goold, C.P., Marek, K.W., and Davis, G.W. (2006). Mechanisms underlying the rapid induction and sustained expression of synaptic homeostasis. Neuron 52, 663–677.

Gervasi, N., Tchenio, P., and Preat, T. (2010). PKA dynamics in a Drosophila learning center: coincidence detection by rutabaga adenylyl cyclase and spatial regulation by dunce phosphodiesterase. Neuron 65, 516–529.

Gratz, S.J., Goel, P., Bruckner, J.J., Hernandez, R.X., Khateeb, K., Macleod, G.T., Dickman, D., and O’Connor-Giles, K.M. (2019). Endogenous Tagging Reveals Differential Regulation of Ca(2+) Channels at Single Active Zones during Presynaptic Homeostatic Potentiation and Depression. J Neurosci 39, 2416–2429.

Handler, A., Graham, T.G.W., Cohn, R., Morantte, I., Siliciano, A.F., Zeng, J., Li, Y., and Ruta, V. (2019). Distinct Dopamine Receptor Pathways Underlie the Temporal Sensitivity of Associative Learning. Cell 178, 60–75 e19.

Hendricks, M., Ha, H., Maffey, N., and Zhang, Y. (2012). Compartmentalized calcium dynamics in a C. elegans interneuron encode head movement. Nature 487, 99–103.

Hige, T., Aso, Y., Modi, M.N., Rubin, G.M., and Turner, G.C. (2015a). Heterosynaptic Plasticity Underlies Aversive Olfactory Learning in Drosophila. Neuron 88, 985–998.

Hige, T., Aso, Y., Rubin, G.M., and Turner, G.C. (2015bb). Plasticity-driven individualization of olfactory coding in mushroom body output neurons. Nature 526, 258–262.

Ichinose, T., Aso, Y., Yamagata, N., Abe, A., Rubin, G.M., and Tanimoto, H. (2015). Reward signal in a recurrent circuit drives appetitive long-term memory formation. Elife 4, e10719.

Inchauspe, C.G., Martini, F.J., Forsythe, I.D., and Uchitel, O.D. (2004). Functional compensation of P/Q by N-type channels blocks short-term plasticity at the calyx of Held presynaptic terminal. J Neurosci 24, 10379–10383.

Ishikawa, T., Kaneko, M., Shin, H.S., and Takahashi, T. (2005). Presynaptic N-type and P/Q-type Ca2+ channels mediating synaptic transmission at the calyx of Held of mice. J Physiol 568, 199–209.

James, T.D., Zwiefelhofer, D.J., and Frank, C.A. (2019). Maintenance of homeostatic plasticity at the Drosophila neuromuscular synapse requires continuous IP3-directed signaling. Elife 8.

Jenett, A., Rubin, G.M., Ngo, T.T., Shepherd, D., Murphy, C., Dionne, H., Pfeiffer, B.D., Cavallaro, A., Hall, D., Jeter, J., et al. (2012). A GAL4-driver line resource for Drosophila neurobiology. Cell Rep 2, 991–1001.

Jing, M., Li, Y., Zeng, J., Huang, P., Skirzewski, M., Kljakic, O., Peng, W., Qian, T., Tan, K., and Wu, R. (2019). An optimized acetylcholine sensor for monitoring in vivo cholinergic activity. bioRxiv, 861690.

Jing, M., Zhang, P., Wang, G., Feng, J., Mesik, L., Zeng, J., Jiang, H., Wang, S., Looby, J.C., Guagliardo, N.A., et al. (2018). A genetically encoded fluorescent acetylcholine indicator for in vitro and in vivo studies. Nat Biotechnol 36, 726–737.

Kim, Y.C., Lee, H.G., and Han, K.A. (2007). D1 dopamine receptor dDA1 is required in the mushroom body neurons for aversive and appetitive learning in Drosophila. J Neurosci 27, 7640–7647.

Konig, C., Khalili, A., Ganesan, M., Nishu, A.P., Garza, A.P., Niewalda, T., Gerber, B., Aso, Y., and Yarali, A. (2018). Reinforcement signaling of punishment versus relief in fruit flies. Learn Mem 25, 247–257.

Liu, C., Placais, P.Y., Yamagata, N., Pfeiffer, B.D., Aso, Y., Friedrich, A.B., Siwanowicz, I., Rubin, G.M., Preat, T., and Tanimoto, H. (2012). A subset of dopamine neurons signals reward for odour memory in Drosophila. Nature 488, 512–516.

Louis, T., Stahl, A., Boto, T., and Tomchik, S.M. (2018). Cyclic AMP-dependent plasticity underlies rapid changes in odor coding associated with reward learning. Proc Natl Acad Sci U S A 115, E448–E457.

Lubbert, M., Goral, R.O., Keine, C., Thomas, C., Guerrero-Given, D., Putzke, T., Satterfield, R., Kamasawa, N., and Young, S.M., Jr. (2019). CaV2.1 alpha1 Subunit Expression Regulates Presynaptic CaV2.1 Abundance and Synaptic Strength at a Central Synapse. Neuron 101, 260–273 e266.

Mao, Z., and Davis, R.L. (2009). Eight different types of dopaminergic neurons innervate the Drosophila mushroom body neuropil: anatomical and physiological heterogeneity. Front Neural Circuits 3, 5.

McGuire, S.E., Le, P.T., and Davis, R.L. (2001). The role of Drosophila mushroom body signaling in olfactory memory. Science 293, 1330–1333.

McGuire, S.E., Le, P.T., Osborn, A.J., Matsumoto, K., and Davis, R.L. (2003). Spatiotemporal rescue of memory dysfunction in Drosophila. Science 302, 1765–1768.

Modi, M.N., Shuai, Y., and Turner, G.C. (2020). The Drosophila Mushroom Body: From Architecture to Algorithm in a Learning Circuit. Annu Rev Neurosci 43, 465–484.

Muller, M., and Davis, G.W. (2012). Transsynaptic control of presynaptic Ca(2)(+) influx achieves homeostatic potentiation of neurotransmitter release. Curr Biol 22, 1102–1108.

Nanou, E., Scheuer, T., and Catterall, W.A. (2016). Calcium sensor regulation of the CaV2.1 Ca2+ channel contributes to long-term potentiation and spatial learning. Proc Natl Acad Sci U S A 113, 13209–13214.

Owald, D., Felsenberg, J., Talbot, C.B., Das, G., Perisse, E., Huetteroth, W., and Waddell, S. (2015). Activity of defined mushroom body output neurons underlies learned olfactory behavior in Drosophila. Neuron 86, 417–427.

Perisse, E., Owald, D., Barnstedt, O., Talbot, C.B., Huetteroth, W., and Waddell, S. (2016). Aversive Learning and Appetitive Motivation Toggle Feed-Forward Inhibition in the Drosophila Mushroom Body. Neuron 90, 1086–1099.

Perkins, L.A., Holderbaum, L., Tao, R., Hu, Y., Sopko, R., McCall, K., Yang-Zhou, D., Flockhart, I., Binari, R., Shim, H.S., et al. (2015). The Transgenic RNAi Project at Harvard Medical School: Resources and Validation. Genetics 201, 843–852.

Pfeiffer, B.D., Ngo, T.T., Hibbard, K.L., Murphy, C., Jenett, A., Truman, J.W., and Rubin, G.M. (2010). Refinement of tools for targeted gene expression in Drosophila. Genetics 186, 735–755.

Placais, P.Y., Trannoy, S., Friedrich, A.B., Tanimoto, H., and Preat, T. (2013). Two pairs of mushroom body efferent neurons are required for appetitive long-term memory retrieval in Drosophila. Cell Rep 5, 769–780.

Rowan, M.J., DelCanto, G., Yu, J.J., Kamasawa, N., and Christie, J.M. (2016). Synapse-Level Determination of Action Potential Duration by K(+) Channel Clustering in Axons. Neuron 91, 370–383.

Sayin, S., De Backer, J.F., Siju, K.P., Wosniack, M.E., Lewis, L.P., Frisch, L.M., Gansen, B., Schlegel, P., Edmondson-Stait, A., Sharifi, N., et al. (2019). A Neural Circuit Arbitrates between Persistence and Withdrawal in Hungry Drosophila. Neuron 104, 544–558 e546.

Scaplen, K.M., Talay, M., Fisher, J.D., Cohn, R., Sorkac, A., Aso, Y., Barnea, G., and Kaun, K.R. (2021). Transsynaptic mapping of Drosophila mushroom body output neurons. Elife 10.

Schroll, C., Riemensperger, T., Bucher, D., Ehmer, J., Voller, T., Erbguth, K., Gerber, B., Hendel, T., Nagel, G., Buchner, E., and Fiala, A. (2006). Light-induced activation of distinct modulatory neurons triggers appetitive or aversive learning in Drosophila larvae. Curr Biol 16, 1741–1747.

Schwaerzel, M., Monastirioti, M., Scholz, H., Friggi-Grelin, F., Birman, S., and Heisenberg, M. (2003). Dopamine and octopamine differentiate between aversive and appetitive olfactory memories in Drosophila. J Neurosci 23, 10495–10502.

Sejourne, J., Placais, P.Y., Aso, Y., Siwanowicz, I., Trannoy, S., Thoma, V., Tedjakumala, S.R., Rubin, G.M., Tchenio, P., Ito, K., et al. (2011). Mushroom body efferent neurons responsible for aversive olfactory memory retrieval in Drosophila. Nat Neurosci 14, 903–910.

Takemura, S.Y., Aso, Y., Hige, T., Wong, A., Lu, Z., Xu, C.S., Rivlin, P.K., Hess, H., Zhao, T., Parag, T., et al. (2017). A connectome of a learning and memory center in the adult Drosophila brain. Elife 6.

Tanaka, N.K., Tanimoto, H., and Ito, K. (2008). Neuronal assemblies of the Drosophila mushroom body. J Comp Neurol 508, 711–755.

Tanimoto, H., Heisenberg, M., and Gerber, B. (2004). Experimental psychology: event timing turns punishment to reward. Nature 430, 983.

Taufiq, A.M., Fujii, S., Yamazaki, Y., Sasaki, H., Kaneko, K., Li, J., Kato, H., and Mikoshiba, K. (2005). Involvement of IP3 receptors in LTP and LTD induction in guinea pig hippocampal CA1 neurons. Learn Mem 12, 594–600.

Tomchik, S.M., and Davis, R.L. (2009). Dynamics of learning-related cAMP signaling and stimulus integration in the Drosophila olfactory pathway. Neuron 64, 510–521.

Tully, T., and Quinn, W.G. (1985). Classical conditioning and retention in normal and mutant Drosophila melanogaster. J Comp Physiol A 157, 263–277.

Xu, C.S., Januszewski, M., Lu, Z., Takemura, S.-y., Hayworth, K., Huang, G., Shinomiya, K., Maitin-Shepard, J., Ackerman, D., and Berg, S. (2020). A Connectome of the Adult Drosophila Central Brain. bioRxiv.

Yamagata, N., Hiroi, M., Kondo, S., Abe, A., and Tanimoto, H. (2016). Suppression of Dopamine Neurons Mediates Reward. PLoS Biol 14, e1002586.

Yamagata, N., Ichinose, T., Aso, Y., Placais, P.Y., Friedrich, A.B., Sima, R.J., Preat, T., Rubin, G.M., and Tanimoto, H. (2015). Distinct dopamine neurons mediate reward signals for short-and long-term memories. Proc Natl Acad Sci U S A 112, 578–583.

Zamponi, G.W., and Currie, K.P. (2013). Regulation of Ca(V)2 calcium channels by G protein coupled receptors. Biochim Biophys Acta 1828, 1629–1643.

Zars, T., Fischer, M., Schulz, R., and Heisenberg, M. (2000). Localization of a short-term memory in Drosophila. Science 288, 672–675.

Zhang, S., and Roman, G. (2013). Presynaptic inhibition of gamma lobe neurons is required for olfactory learning in Drosophila. Curr Biol 23, 2519–2527.

Zhang, X., Noyes, N.C., Zeng, J., Li, Y., and Davis, R.L. (2019). Aversive Training Induces Both Presynaptic and Postsynaptic Suppression in Drosophila. J Neurosci 39, 9164–9172.

Zhao, X., Lenek, D., Dag, U., Dickson, B.J., and Keleman, K. (2018). Persistent activity in a recurrent circuit underlies courtship memory in Drosophila. Elife 7.

